# Functional Adaptations of Endogenous Retroviruses to the *Drosophila* Host Underlie their Evolutionary Diversification

**DOI:** 10.1101/2023.08.03.551782

**Authors:** Kirsten-Andre Senti, Dominik Handler, Baptiste Rafanel, Carolin Kosiol, Christian Schlötterer, Julius Brennecke

**Affiliations:** Institute of Molecular Biotechnology of the Austrian Academy of Sciences (IMBA), Vienna BioCenter (VBC); Dr. Bohr-Gasse 3, 1030 Vienna, Austria; Vienna BioCenter PhD Program, Doctoral School of the University of Vienna and Medical University of Vienna, Vienna, Austria; University of St Andrews, Centre for Biological Diversity; St Andrews, Scotland, UK; Institut für Populationsgenetik, Vetmeduni Vienna; Veterinärplatz 1, 1210 Vienna, Austria

## Abstract

Transposable elements profoundly affect the biology and evolution of their hosts, yet their own evolutionary dynamics remain poorly understood. Here, we investigate insect endogenous retroviruses (iERVs), a monophyletic group of LTR retrotransposons that have acquired the trait of infectivity, likely through capture of a Baculovirus *envelope* gene. In *Drosophila* ovaries, iERVs with functional *envelope* have adapted their *cis*-regulatory sequences to be expressed in any somatic cell type, from where they infect the germline. Strikingly, related retroviruses show distinct expression patterns, indicating niche partitioning. In contrast, all non-infectious iERVs that emerged through secondary *envelope*-loss are specifically expressed in the germline. Co-evolving with iERVs, the genome-protecting piRNA pathway has assimilated iERV promoter and sequence information into piRNA clusters, underscoring the functional significance of iERV expression in somatic niches. We propose that the evolutionary innovation of cell-to-cell infectivity has triggered the adaptive radiation of iERVs through trait diversification and antagonistic virus-host interactions, processes that likely underpin niche-specific expression of endogenous retroviruses in vertebrates as well.

## Introduction

The relationship between transposable elements (TEs) and their host organisms is, at its core, a parasite-host conflict. TEs exploit the cellular resources of their hosts to generate new insertions, compromising the integrity of the host genome and resulting in significant fitness costs (*1*). Consequently, host organisms have evolved diverse strategies to mitigate the deleterious effects of TEs (*2, 3*). This dynamic has led to a relentless genetic conflict, with hosts employing mechanisms to detect and suppress TEs, while TEs strive to evade these restriction systems and replicate within the host genome.

The emergence of genome defense systems in eukaryotes has led to the tolerance of TEs (*4*), resulting in their accumulation within host genomes (*5, 6*). This abundance of TE-derived sequences has profound implications for genome structure and chromosome biology (*7–9*). The co-evolution between TEs and their hosts is also a major driver of biological innovation (*4, 10–12*). For instance, many evolutionary leaps such as the emergence of gene regulatory networks, adaptive immunity in vertebrates, or placental development in mammals are linked to the TE-host conflict through the co-option of TE-derived regulatory or coding sequences (*13–15*). However, while we increasingly appreciate the profound impact of TEs on host evolution, our knowledge of the evolutionary dynamics underlying the diversification of TEs themselves remains limited, foremost due to a lack of suitable study systems.

*gypsy*/*Ty3* elements (the *Metaviridae*) are an ancient class of long terminal repeat (LTR) retrotransposons found in plants, fungi, and animals (*16, 17*). Most *Metaviridae* contain two open-reading frames, *gag* and *pol*, which encode the capsid proteins and enzymes for replication and genome integration, respectively. In animals, the replication of *Metaviridae* occurs within germline cells, but is repressed by the PIWI interacting RNA (piRNA) pathway (*18, 19*). Intriguingly, during evolution some *Metaviridae* lineages, the Errantiviruses or insect endogenous retroviruses (iERVs), have acquired a key biological novelty: infectivity (*17, 20*). This evolutionary leap was based on the capture of a third open reading frame related to F-type Envelope proteins of DNA baculoviruses (*21, 22*). F-type Envelope proteins facilitate the fusion of viral particles with host cell membranes, and acquisition of *env* therefore transformed an intracellularly replicating LTR retroelement into an infectious retrovirus. The resulting iERVs exhibit a highly similar architecture compared to the phylogenetically separate basal retroviruses found in vertebrates (e.g., alpha clade of *Retroviridae*). However, unlike the *Retroviridae*, iERVs did not adopt a lifestyle as horizontally transmitted, exogenous viruses. Instead, the experimentally studied iERVs, *gypsy* and *ZAM*, are expressed in follicle cells of *Drosophila* ovaries defective in the piRNA pathway and infect the adjacent oocyte as viral particles to integrate new copies of themselves into the germline genome (*23–26*). This distinctive pattern suggests that iERVs have evolved a sophisticated vertical transmission strategy by leveraging neighboring somatic cells to infect the germline, thereby maintaining their presence in the host genome.

Here, we systematically investigate the evolution and biology of the iERV clade within the context of the *Drosophila melanogaster* host. Our study unveils a direct correlation between the trait of infectivity and the colonization of the entire ovarian soma by iERVs, establishing it as prominent niche for their replication. During this process, iERVs have diversified greatly, in part by adapting their *cis*-regulatory elements to the multiple somatic cell types. Intriguingly, the secondary loss of the *envelope* gene has repeatedly resulted in the emergence of non-infectious iERVs, all of which have switched their expression patterns to the germline. Our study further reveals the co-evolutionary dynamics between iERVs and the piRNA pathway, elucidating how tissue-specific regulation of piRNA source loci has adaptively evolved in response to iERV radiation. Thus, iERVs emerge as a paradigm for elucidating the intricate evolutionary dynamics of TE-host interactions in animal gonads.

## Results

### iERVs constitute a monophyletic LTR clade comprised of retroviruses and derived retroelements

The *Drosophila melanogaster* genome harbors several dozen LTR retrotransposon lineages, defined by distinct consensus sequences, belonging to the *Belpaoviridae* and the *Metaviridae* (*27, 28*). We systematically curated and annotated the main open reading frames (ORFs) for forty-nine lineages and estimated a phylogenetic tree based on a protein sequence alignment for full length *pol*, the most conserved gene in LTR retrotransposons. This analysis divided the *Metaviridae* into the five known clades, *gypsy/mdg3, gypsy/osvaldo, gypsy/mdg1, gypsy/chimpo*, and *gypsy/gypsy* (Fig. 1A) (*27*).

**Fig. 1:**
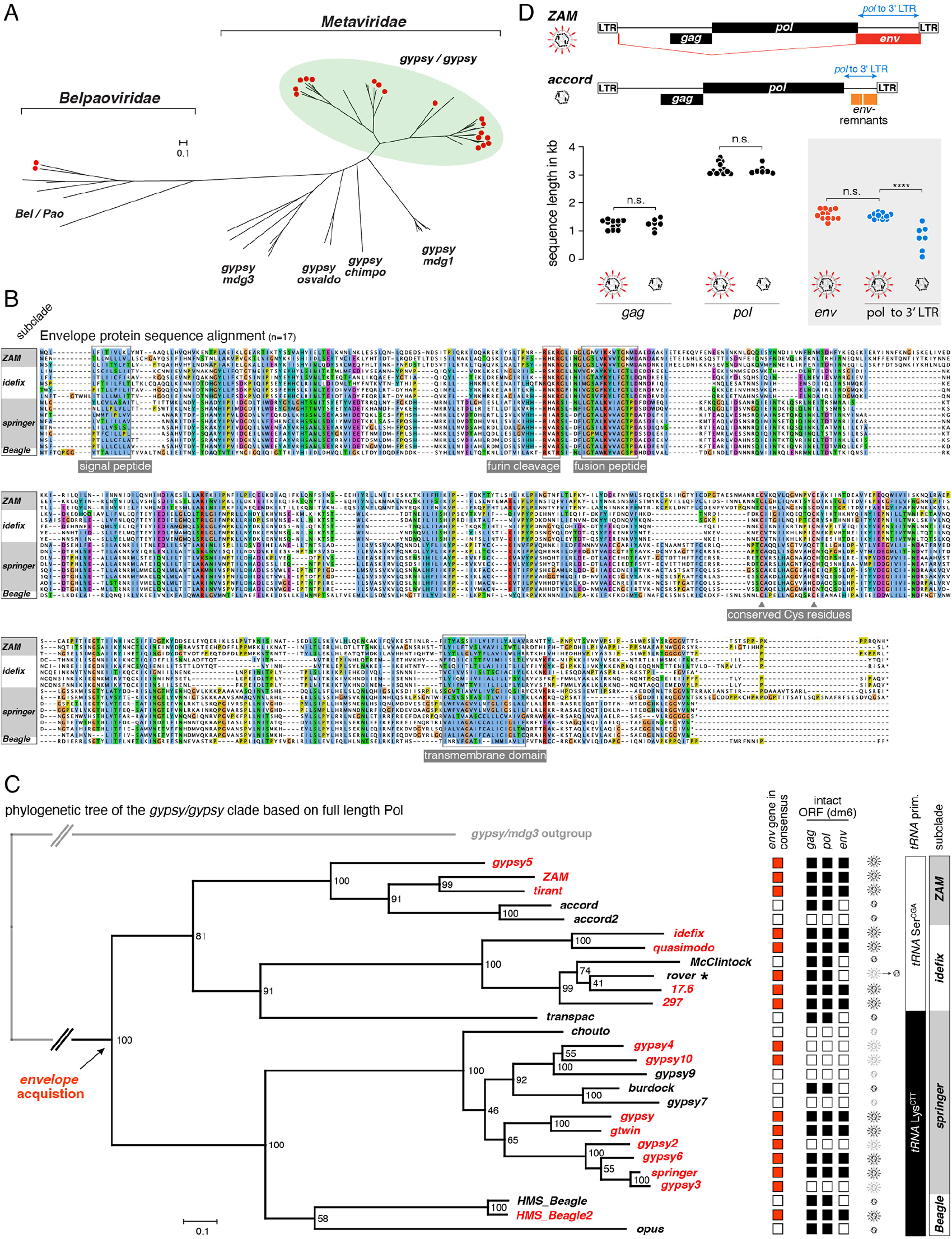
Envelope acquisition and diversification of insect endogenous retroviruses in *Drosophila*. **(A)** Phylogenetic tree, based on full length Pol of all *Belpaoviridae* and *Metaviridae* LTR-retroelement consensus sequences from *Drosophila melanogaster*. LTR-element subclades are indicated and retroviral species with a full-length *env-F* gene are marked with a red dot. **(B)** Alignment of all seventeen full-length iERV Env-F protein consensus sequences. From top to bottom Env sequences are from *gypsy5, ZAM, tirant, idefix, quasimodo, rover, 297, 17.6, gypsy4, gypsy10, gypsy, gtwin, gypsy2, gypsy6, springer, gypsy3, HMS-Beagle 2*. Subclades and conserved, functional protein features are indicated. **(C)** Phylogenetic tree, based on full length Pol, of all iERVs from *Drosophila melanogaster* (outgroup: *gypsy*/*mdg3* clade; numbers indicate bootstrap values). Lineages with full-length *env-F* are indicated in red, retroelement revertants in black. To the right, the inferred activity status of each lineage based on intact *gag*, *pol*, *env-F* open reading frames in at least one insertion in *dm6* is shown. Retroviral and retroelement (capsid only) symbols indicate active (black) or inactive (grey) species (the asterisk labels the transition element *rover*). **(D)** Cartoon depicting the *ZAM* retrovirus (top) and *accord* retroelement revertant with *env-F* fragments (bottom). The jitter plot shows the length (in nucleotides) of *gag*, *pol*, *env-F* open reading frames as well as the *pol* to 3’ LTR distance for all active iERVs (statistical significance based on unpaired, non-parametric Mann-Whitney test).

Nineteen LTR retrotransposon lineages contain a third ORF, a predicted *envelope* gene, downstream of *gag* and *pol* (Fig. 1A; red dots). Except for *roo* and *rooA,* all these lineages belong to the *gypsy/gypsy* elements, forming the most diverse *Metaviridae* clade, also known as *Errantiviruses* or insect endogenous retroviruses (iERVs) (*17, 20, 29*). As Envelope proteins are translated from spliced subgenomic transcripts (*23, 25, 30–32*), we experimentally mapped the splice junctions for all active lineages and predicted those of the inactive ones (Suppl. Fig. 1 and 2). This revealed that the main *env* ORF is spliced to a short peptide upstream of *gag* in the *ZAM* and *Beagle* subclades, to an AU dinucleotide (part of the methionine start codon) in the *springer* subclade, or to a Gag peptide in the *idefix* subclade. A sequence alignment of the resulting seventeen full-length Envelope proteins demonstrated strong similarities (Fig. 1B) and, in agreement with prior studies, clear relationships to F-type Envelope glycoproteins found in baculoviruses (Suppl. Fig. 3) (*21, 22*). These similarities include highly conserved sequence features, encompassing an N-terminal signal peptide, a furin cleavage site, a fusion peptide, conserved cysteine residues for the formation of stabilizing di-sulfide bonds, and a C-terminal transmembrane domain (Fig. 1B) (*33, 34*). We refer to the third ORF in iERVs as *env-F*.

**Fig. 2:**
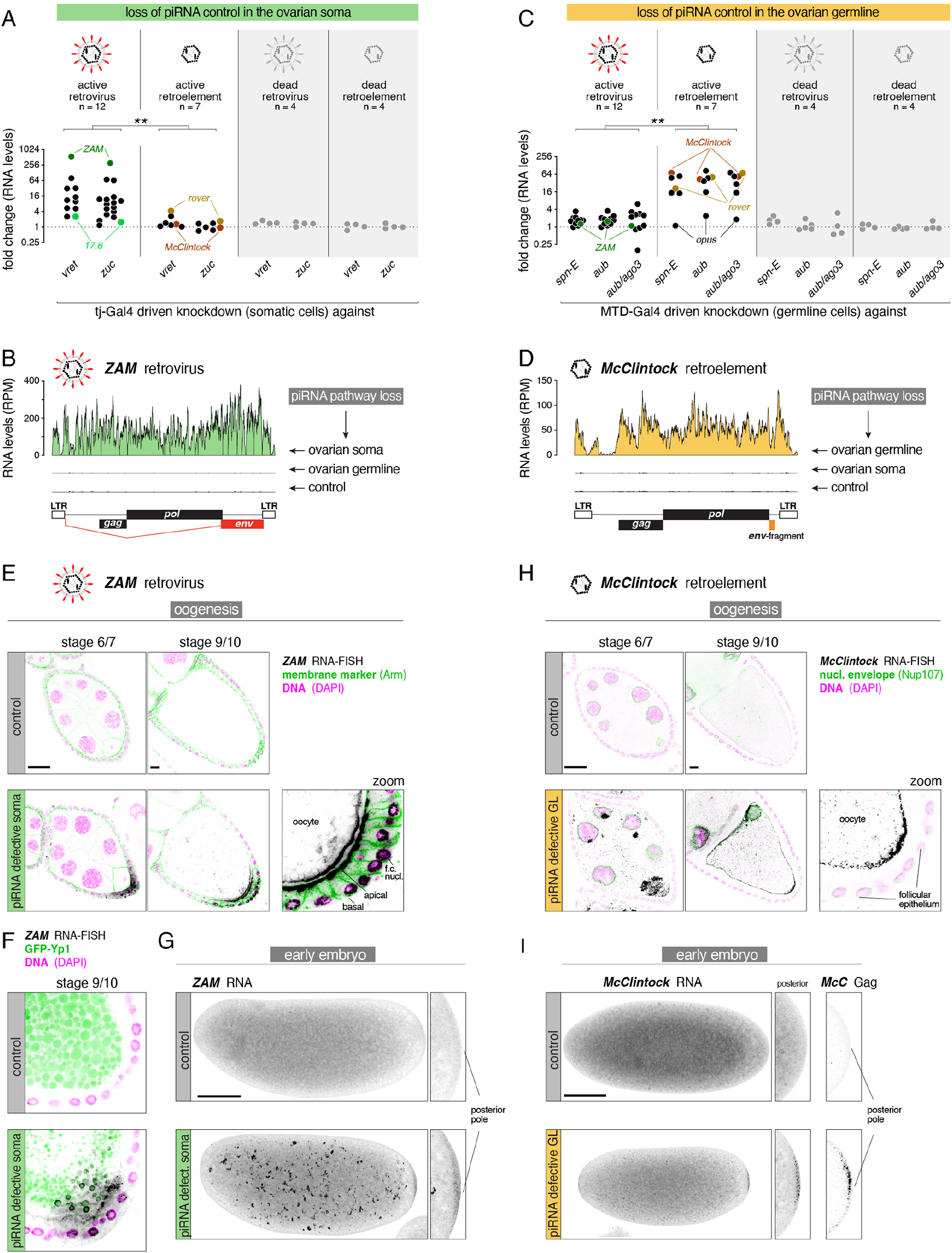
Replication strategies of infectious versus non-infectious iERVs. **(A, C)** Shown are fold changes in ovarian steady state levels of iERV transcripts, grouped into active and inactive retroviruses and retroelements, upon loss of somatic piRNA pathway control (A) (*tj*-Gal4 driven long dsRNA hairpins against *vreteno* or *zucchini*) or loss of germline piRNA pathway control (C) (*MTD*-Gal4 driven shRNAs against *spn-E*, *aub*, or *aub*+*ago3*). The *tirant* retrovirus cannot be conclusively analyzed as it is not present in most strains used in this study. **(B, D)** Shown are normalized transcript levels (RPM) of the representative retrovirus *ZAM* (B) and the retroelement *McClintock* (D) in control ovaries or ovaries lacking somatic or germline piRNA pathway control (genotypes: *tj*-Gal4 > *vreteno^GD^*, *MTD*-Gal4 > *aub+ago3* sh, *tj*-Gal4 > *arrestin2^GD^* in panel B; and *MTD*-Gal4 > *aub+ago3* sh, *tj*-Gal4 > *vreteno^GD^*, *MTD*-Gal4 > *white* sh in panel D). **(E)** RNA-smFISH based expression analysis for the retrovirus *ZAM* in egg chambers of indicated stage from control ovaries and from ovaries with defective somatic piRNA pathway (*tj*-Gal4 driven dsRNA hairpin against *vreteno* or against *arrestin2* for control; error bars: 20µm; cell outlines were stained with an anti-Armadillo antibody and DNA with DAPI). The magnified panel shows the posterior pole of the growing stage 9/10 oocyte with adjacent follicle cells (f.c.). **(F)** As in (E) but with yolk granules labelled with YP1-GFP in green. **(G)** As in (E) but with RNA-smFISH against the *McClintock* retroelement and in ovaries with defective germline piRNA pathway (*MTD*-Gal4 driven shRNA against *aub*+*ago3* or against *white* for control). **(H, I)** RNA-smFISH based expression analysis for *ZAM* (G) or *McClintock* (H) in pre-blastoderm embryos with less than 32 nuclei laid by females lacking somatic (F) or germline (H) piRNA pathway control (genotypes: *tj*-Gal4 > *vreteno^GD^* or *arrestin2^GD^*in panel F, and *MTD*-Gal4 driven shRNA against *aub* or *white* in panel G; images show maximum intensity Z projections; scale bars: 100µm). Magnified panels show accumulation of FISH signal at the posterior pole (also shown with an antibody staining against *McClintock* Gag in panel H).

With a complete set of curated iERV consensus sequences at hand, we performed a systematic phylogenetic analysis. Independent trees estimated from sequence alignments of both full-length Pol and the faster diverging Gag core domain, using two independent maximum likelihood methods, showed robust and congruent phylogenetic architectures (Fig. 1C; Suppl. Fig. 4A, B). A phylogenetic tree estimated from an alignment of the seventeen complete Env-F sequences mirrored the Pol and Gag-trees, arguing against independent *env-F* gain events or *env-F* exchange through retroviral recombination among iERVs (Suppl. Fig. 4 C). Notably, eleven iERV lineages across different subclades lack a full length *env-F* open reading frame (Fig. 1C). We found that these lineages harbor sequence stretches of variable length between the end of *pol* and their 3’ LTR (Fig. 1D). In almost all cases, these sequences contain mutated yet discernible *env-F* fragments (Suppl. Fig. 4D, 5). We infer that all eleven iERV lineages lacking functional *env-F* evolved from infectious retroviruses through *env-F* sequence degradation.

To gain insights into the evolutionary dynamics of *env-F*, we utilized the binary character state speciation and extinction model (BiSSE) (*35*). We estimated a likelihood of 99.1% that a retrovirus with functional *env-F* gene was at the base of the iERV clade (Suppl. Fig. 6A). Furthermore, a Bayesian analysis using functional *env-F* as a binary trait character revealed that retroviruses diversified at a higher rate than retroelements (Suppl. Fig. 6B), although this difference did not reach statistical significance due to the limited number of iERV lineages present in *Drosophila melanogaster* for the analysis (*36*).

We finally considered the *env-F* status along with the integrity of *gag* and *pol* ORFs in all dm6 insertions, and classified iERV lineages as either *env-F*-encoding retroviruses (12 active, 4 inactive) or *env-F*-deficient retroelements (7 active, 4 inactive) (Fig. 1C). Consistent with their annotated status, insertions of inactive retroviruses and retroelements primarily localize within heterochromatic regions and exhibit substantial pairwise LTR divergence (Suppl. Fig. 6C). We categorized *rover* as an active retroelement undergoing a transition from retrovirus to retroelement: four *rover* insertions contain a full-length *env-F* but defective *gag* or *pol* ORFs (inactive retroviruses), whereas eight almost identical insertions harbor intact *gag* and *pol* ORFs but defective *env-F* (active retroelements). In conclusion, our findings provide robust support for iERVs being a monophyletic LTR retrotransposon clade, originating from an *env-F* encoding ancestral retrovirus, and comprising numerous active retroviral and derived, non-infectious retroelement lineages.

### Infectious and non-infectious iERVs evolved distinct expression and replication strategies

Considering the closely related retrovirus and derived retroelement lineages among iERVs, we examined the potential relationship between their trait of infectivity (characterized by the presence of a functional *env-F*) and their expression. Our primary focus was on the ovary of adult flies, where germline and somatic cells maintain direct contact during the entire oogenesis process (*37, 38*).

In wildtype flies, distinct piRNA pathways silence TEs in the ovarian soma and germline, respectively (*18, 39*). To systematically reveal the evolved spatio-temporal expression patterns of iERVs, we employed tissue-specific, transgenic RNA interference (*40–43*). We used *tj*-Gal4 to deplete the essential piRNA pathway components *vreteno* or *zucchini* in all ovarian somatic cells (soma-knockdown) and MTD-Gal4 to deplete the essential components *spindle-E, aubergine* or *aubergine/ago3* in all ovarian germline cells (germline-knockdown) (Suppl. Fig. 7). We monitored transcript levels by polyA RNA-seq. Intriguingly, active retroviruses were consistently de-repressed in soma-knockdown ovaries but not in germline-knockdown ovaries (Fig. 2A, B; Suppl. Fig. 8A), whereas active retroelements were exclusively derepressed in germline-knockdown ovaries (Fig. 2C, D; Suppl. Fig. 8B). The steady-state RNA levels of inactive iERV lineages, represented only by defective insertions, remained unaffected in both conditions. These findings strongly suggest that retroviruses and retrovirus-derived retroelements are generally expressed in fundamentally opposing tissues of the ovary (soma versus germline).

Irrespective of the tissue in which they are transcribed, iERVs must integrate new copies of themselves into the host germline genome for their survival. To further investigate iERV replication strategies, we employed highly sensitive single molecule fluorescent RNA in situ hybridization (smFISH) (*43, 44*). We focused on two representative iERVs, the retrovirus *ZAM* and the retro-element *McClintock*. In control ovaries and in germline knockdown ovaries, *ZAM* transcripts were undetectable. In contrast, *ZAM* exhibited strong expression in soma-knockdown ovaries, reaching maximum levels in stage 9/10 egg chambers (Fig. 2E; Suppl. Fig. 9A) (*25*). *ZAM* transcripts were enriched in nuclei and at the apical membrane of posterior follicle cells, as well as at the adjacent oocyte membrane and within the oocyte cytoplasm, despite the complete absence of *ZAM* expression in the germline. Within the oocyte, *ZAM* transcripts localized to yolk granules labelled by GFP-tagged Yolk Protein 1 (Fig. 2F) (*45, 46*), supporting the prior notion that the *ZAM* retrovirus highjacks the vitellogenesis pathway for its transfer from follicle cells into the late-stage oocyte (*26*). To quantify germline transmission directly, we sequenced poly-adenylated RNA from dechorionated 0-1h old embryos, which solely contain maternally provided germline RNAs. In embryos laid by mothers with defective somatic piRNA control, *ZAM* transcript levels were 27-fold higher than in control embryos (Suppl. Fig. 8C, 9B). In these embryos, *ZAM* RNA was found in disintegrating yolk granules and at the posterior pole (Fig. 2G). We also observed soma-to-oocyte transfer for the retroviruses *springer* and *gypsy* (Suppl. Fig. 8C, 9B-E).

In stark contrast to *ZAM*, transcripts of the non-infectious retroelement *McClintock* were exclusively observed in germline knockdown ovaries (Fig. 2C, D; Suppl. Fig. 8A, B, 9G). *McClintock* transcripts and the capsid protein Gag were found in nurse cells and enriched in the transcriptionally inactive oocyte, indicative of transport of *McClintock* capsids from nurse cells through ring-canals (Fig. 2H; Suppl. Fig. 9H) (*43, 47*). *McClintock* transcript levels were increased ∼150-fold in 0-1h old embryos laid by mothers with germline knockdown compared to control embryos (Suppl. Fig. 8D, 9F). Similar to *ZAM*, *McClintock* transcripts, as well as transcripts from other retroelements such as *burdock* and *rover,* accumulated at the posterior pole of embryos (Fig. 2I; Suppl. Fig, 9I, J). Thus, despite being transcribed in entirely different tissues, both *McClintock* and *ZAM* target their genetic material to the posterior pole plasm, where the primordial germ cells of the embryo will form.

### Closely related retroviruses display niche partitioning in the ovarian soma

The transcript levels of infectious retroviruses increased to very different extents in soma-knockdown ovaries, ranging from 3-fold (*17.6*) to almost 600-fold (*ZAM*) (Suppl. Fig. 8A). To investigate whether this variability was associated with temporally or spatially restricted expression domains, we performed RNA smFISH for all active retroviruses for the entire oogenesis process. Remarkably, this revealed that each retroviral lineage, while repressed in control ovaries, exhibits a distinct expression pattern in soma-knockdown ovaries, and that collectively retroviruses occupy essentially every cell type of the ovarian soma (Fig. 3A summarizes the detailed expression data compiled in Suppl. Fig. 10).

**Fig. 3:**
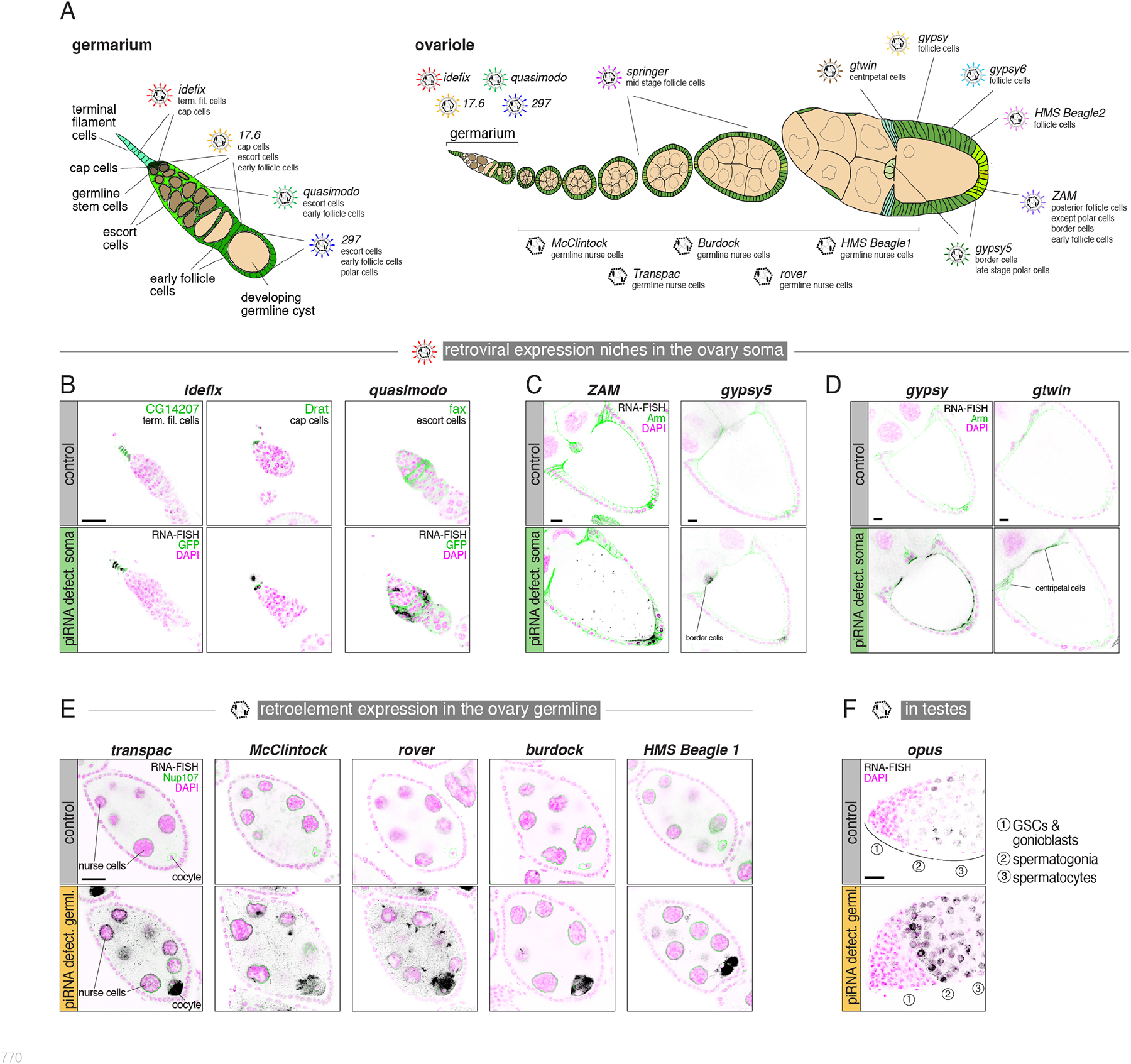
Gonadal expression niches of infective and non-infective iERVs. **(A)** Cartoon summarizing the major iERV expression domains in the germarium (left) and entire ovariole (right). Germline cells are shown in beige, somatic cells in green. **(B-D)** RNA-smFISH based expression analysis of closely related retroviral pairs in control ovaries or ovaries lacking somatic piRNA pathway control (*tj*-Gal4 driven dsRNA hairpins against *arrestin2* or *vreteno*). Panel B shows *idefix* and *quasimodo* (black) in the germarium with relevant cell types identified by indicated GFP-trap lines in green and DAPI in magenta. Panels C and D show *ZAM* and *gypsy5* (C) and *gypsy* and *gtwin* (D) expression (black) in stage 9/10 egg chambers with cell outlines marked by anti-Armadillo in green and DAPI in magenta. *gypsy5* images in panel C show maximum intensity projections of five Z-sections. **(E)** RNA-smFISH based expression analysis of five retroelement revertants in stage 6/7 egg chambers of control ovaries or of ovaries lacking germline piRNA control (*MTD*-Gal4 driven shRNAs against *white* or *aub*+*ago3*; DAPI is shown in magenta, GFP-Nup107 in green and indicated smFISH in black). **(F)** RNA-smFISH based expression analysis of *opus* (black) in the anterior tip of testes lacking germline piRNAs compared to controls (*nanos*+*bam*-Gal4 driven shRNAs against *aub*+*ago3* or *white* as control; DAPI is shown in magenta; major germline cell types are indicated; scale bar: 20µm).

Retroviruses that exhibited a strong de-repression at the ovary RNA-seq level (*ZAM*, *gypsy*, *springer*, *gypsy6, HMS-Beagle 2*) were expressed in broad domains, while those with moderate de-repression (*idefix*, *17.6*, *quasimodo*, *297*, *gypsy5*) had restricted expression niches (Suppl. Fig. 8A, 10). We infer that retroviruses are robustly transcribed within their cognate niches. Moreover, viruses from the *idefix* subclade were expressed during early stages, whereas those belonging to the *ZAM* and *springer* subclades were predominantly expressed at late stages (Fig. 3A). These findings suggest that viruses from different iERV subclades may target the germline genome at distinct developmental stages of oogenesis.

Notably, closely related retroviral lineages sharing a recent common ancestor displayed spatially close but distinct expression patterns in every case examined. For instance, the *idefix*/*quasimodo* pair showed expression in non-overlapping cell populations of the germarium, with *idefix* being transcribed in terminal filament and cap cells, and *quasimodo* in escort cells (Fig. 3B). Similarly, the retroviruses *17.6* and *297* were detected in cap cells, escort cells and early follicle cells (*17.6*) or in escort cells, early follicle cells and polar cells (*297*) (Suppl. Fig. 11A, B). Similar patterns were observed for related retroviruses expressed during late oogenesis stages: *ZAM* was highly expressed in posterior follicle cells, whereas *gypsy5* was mainly expressed in border cells and polar cells (Fig. 3C). Likewise, *gypsy* was expressed in all main body follicle cells of late-stage egg chambers, while the related *gtwin* virus was expressed in centripetal cells (Fig. 3D). Lastly, *springer* and *gypsy6* were expressed in overlapping pattern in main body follicle cells, but initiated expression at distinct developmental stages (stage 6/7 versus stage 9/10, respectively) (Suppl. Fig. 11C, D). In conclusion, *env-F* encoding iERVs have collectively occupied literally all somatic cell types in the ovary, but individual lineages are expressed in distinct, yet often overlapping cell-type niches (Fig. 3A).

### iERVs that lost env-F, adapted their expression back to the germline

During iERV diversification, functional *env-F* has been lost multiple times, leading to the emergence of eleven retroelement lineages, of which seven are active (*accord*, *rover*, *McClintock*, *transpac*, *burdock*, *HMS-Beagle*, *opus*) (Fig. 1C; Suppl. Fig. 6C). Based on RNA-seq experiments, retroelement lineages were strongly de-repressed in germline knockdown ovaries but not in soma knockdown ovaries (Suppl. Fig. 8A, B). Notable exceptions were the transition element *rover*, which was also weakly de-repressed in the soma, and *opus* which was not de-repressed in either knockdown. RNA smFISH experiments validated these findings, supporting the exclusive expression of retroelements in germline cells, ultimately resulting in the pronounced accumulation of TE transcripts in the maturing oocyte (Fig. 3E; Suppl. Fig. 10, 12). In all examined cases, expression commenced in differentiating cystoblasts and persisted until late-stage egg chambers (Suppl. Fig. 13). We hypothesized that the outlier *opus* might be expressed in testes instead of ovaries. Indeed, smFISH experiments revealed that *opus* was strongly de-repressed in spermatogonia lacking piRNA control (Fig. 3F). These findings collectively suggest that while infectious retroviruses diversified their expression to different niches in the ovarian soma, the derived non-infectious retroelements converged their expression to germline cells. A remarkable example of this expression switch is the pair of retrovirus *HMS Beagle-2* and retroelement *HMS Beagle*, which share a sequence identity of over 90% for Gag-core and Pol. In piRNA-deficient ovaries, *HMS Beagle-2,* containing a complete *env-F* gene, exhibited exclusive expression in the soma, whereas *HMS Beagle*, lacking *env-F*, was expressed solely in the germline (Suppl. Fig. 10, 12).

### iERV expression niches are defined by intrinsic cis-regulatory sequences

The specific expression patterns exhibited by different iERVs may arise from distinct regulatory sequences that are within the TE itself or from the host genome nearby individual iERV insertions. We found that LTR and 5’ UTR sequences, which typically harbor the transcription regulatory elements in LTR retroviruses and retroelements (*48*), are the most divergent segments of iERVs (Fig. 4A; Suppl. Fig. 14). Their only conserved sequence motifs are at the LTR boundaries (presumably integrase recognition sites) and at the beginning of the 5’ UTR where the primer binding sites for specific tRNAs reside (*20*). These findings suggest that the pronounced cell type-specific transcription patterns of the different iERV lineages could arise from sequence variations in their non-coding regions.

**Fig. 4:**
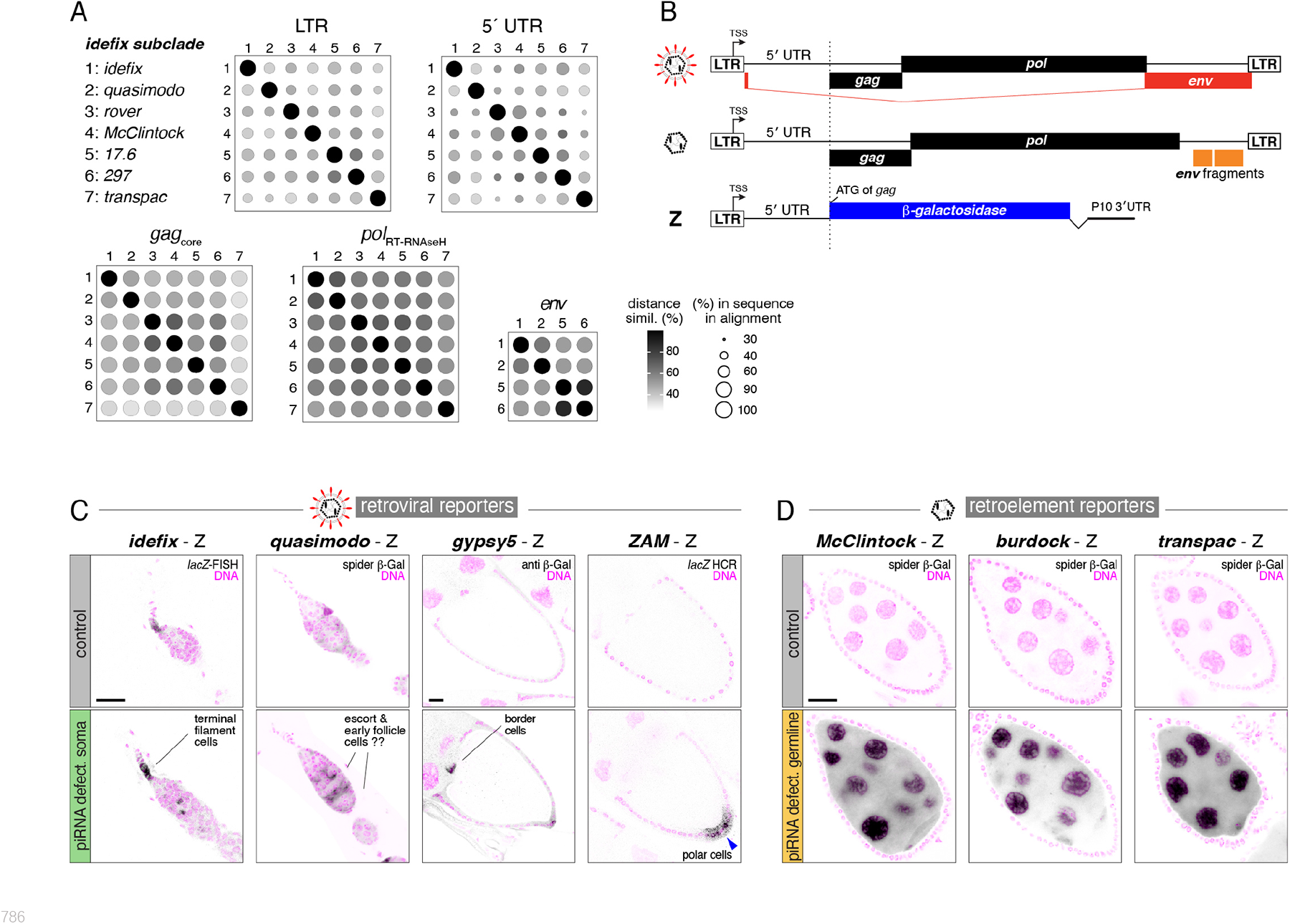
*cis*-regulatory sequences within iERVs define their expression patterns. **(A)** Shown are pairwise similarities between the DNA sequences for LTR, 5’ UTR, the protein domains Gag core, RT-RNAseH, and full length spliced Env-F. Circle size indicates percentage of nucleotides in aligned positions, greyscale indicates pairwise identity in percent, calculated in Clustal Omega without correction among *idefix* subclade species (see Supp. Fig. 14 for other subclades). **(B)** Cartoon depicting the architecture of retroviruses, retroelements, and the lacZ-reporter transgenes tested in C and D for *quasimodo, gypsy5*, *McClintock, burdock*, and *transpac*. The *idefix* and *ZAM* reporters are previously published LTR-lacZ fusions. **(C, D)** Analysis of retroviral (C) or retroelement (D) lacZ reporter expression in control ovaries and in ovaries lacking somatic or germline piRNA pathway control (genotypes are *tj*-Gal4 driven dsRNA hairpins against *vreteno* or *arrestin2* as control in panel C, and *MTD*-Gal4 driven shRNAs against *aub* or *white* as control in panel D; germarium stages are shown for *idefix* and *Quasimodo*, stage 9/10 egg chambers for *gypsy5* and *ZAM*, and stage 6/7 egg chambers for retroelements; scale bars: 20µm).

To test this hypothesis, we examined published lacZ reporter fly lines for the retroviruses *idefix* and *ZAM* (*30, 49*), and generated new reporter lines for the retroviruses *quasimodo* and *gypsy5*, as well as for the retroelements *burdock*, *McClintock*, and *transpac* (Fig. 4B). All newly generated reporter transgenes were inserted into the same genomic locus lacking transcriptional activity in ovaries. In wildtype ovaries, all reporters showed no or only very weak expression, presumably due to stringent piRNA-mediated repression. All retroviral reporters were exclusively expressed in soma knockdown ovaries, mirroring the corresponding endogenous retroviruses (Fig. 4C). In contrast, all retroelement reporters were only expressed in germline knockdown ovaries, again mirroring the expression of their endogenous counterparts (Fig. 4D). We conclude that individual retroviral lineages have adapted their *cis*-regulatory elements (CREs) to drive expression in distinct somatic cell types. Whenever a lineage lost its infectivity trait, it also lost its soma specific CREs but gained CREs that facilitate transcription in the germline.

### piRNA clusters co-evolve with iERVs through the co-option of retroviral sequences

Two distinct piRNA pathways are active in the soma and germline of the ovary, utilizing separate genomic piRNA source loci (*18, 39*). In germline cells, most TE-targeting piRNAs are derived from heterochromatic loci that are specified by the HP1-variant protein Rhino and transcribed in a non-canonical manner on both strands (*50–52*). Consistent with this, ovaries with germline-specific depletion of *rhino* displayed strongly reduced levels of piRNAs antisense to the active, germline-expressed retroelements among iERVs (Suppl. Fig. 15A). The only exception was the transition element *rover* (see below).

Despite their Rhino-dependency, we found no corresponding retroelement insertion or silencing potential (determined as 25mer sequences in cluster loci mapping to TE sequences) in the major Rhino-dependent piRNA clusters *42AB*, *38C*, *80F* (Suppl. Fig. 15B). Also, retroelement piRNAs were unaffected in flies lacking piRNA clusters *38C* and *42AB* (Suppl. Fig. 15C) (*53*). Instead, several stand-alone insertions of retroelements within local heterochromatin domains bound by Rhino and Kipferl acted as piRNA source loci as evidenced by piRNAs mapping to flanking genome-unique sequences (Suppl. Fig. 15D) (*52, 54–56*). These findings highlight the adaptive capacity of the germline piRNA system, where TE control does not necessarily depend on capturing newly invading TEs within existing large piRNA clusters but involves the Rhino-dependent conversion of individual TE insertions into piRNA source loci.

Rhino is not expressed in somatic cells of the ovary, where piRNA clusters are canonical RNA polymerase II transcription units with defined promoters (*52, 57*). The silencing spectrum of the somatic piRNA pathway relies on TE insertions within a piRNA cluster, specifically in the antisense orientation to the cluster’s unidirectional transcription. The primary piRNA cluster in the soma, *flamenco*, is a several hundred kilobase long locus on the X-chromosome that is transcribed from a single promoter. Retroviral insertions are enriched in *flamenco* (*39, 57*), and flies carrying specific *flamenco* permissive alleles, exhibit de-repression of the *gypsy*, *ZAM* and *idefix* retroviruses (*58–60*). Thus, a single locus, *flamenco*, might be responsible for silencing the entire spectrum of retroviruses throughout the ovarian soma.

To test the “single cluster” model, we first analyzed *flamenco*’s expression. smFISH experiments revealed that *flamenco* is transcribed in all somatic cell types of the ovary, except for terminal filament cells (Fig. 5A; Suppl. Fig. 7C). We then analyzed *flamenco*’s sequence composition in relation to the iERV phylogeny. To account for the sequence decay of TE fragments, we determined the percentage of full-length iERV sequences found within *flamenco* in antisense orientation as 25-mer sequences. This analysis revealed a striking bias in *flamenco*’s TE content: While active and inactive retroviruses are strongly represented (*61*), none of the active or inactive iERVs that had transitioned to retroelements is present in *flamenco* (Fig. 5B). The sole exception was once more the retroelement *rover*, whose insertion in *flamenco* contains an intact *env-F* gene, strongly supporting its recent status as infectious retrovirus.

**Fig. 5:**
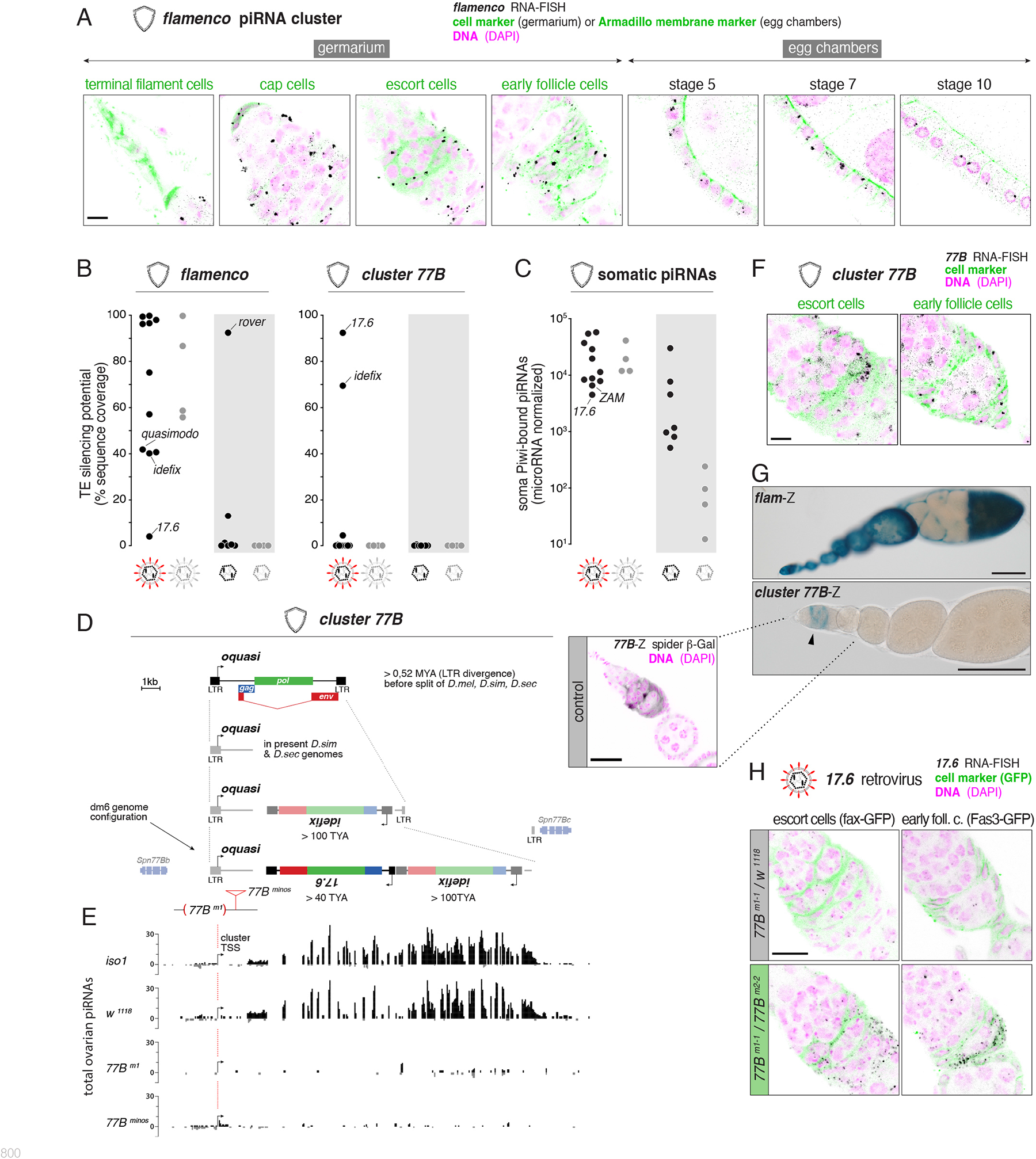
Co-evolution of the somatic piRNA pathway with infectious iERVs. **(A)** Detailed RNA-smFISH based expression analysis of the somatic *flamenco* piRNA cluster (black) in ovarian somatic cells (GFP-trap lines in green indicate cell types in the germarium; for later stages, Armadillo labels cell outlines; DAPI is shown in magenta, oogenesis stages are indicated; scale bars: 10µm.) **(B)** Shown is the silencing potential within indicated somatic piRNA clusters against active and inactive iERVs (retroviruses and retroelements) as fraction of TE sequence found as 25mers (zero mismatches) in antisense orientation in the cluster. **(C)** Shown are microRNA-normalized levels of somatic, Piwi-bound piRNAs antisense to active and inactive retroviruses and retroelements among the iERVs (zero mismatches). **(D)** Proposed evolutionary trajectory of the *D. melanogaster* specific *77B* piRNA cluster that originated from an old *quasimodo* retrovirus (*oquasi*) insertion between the *Spn77Bb* and *Spn77Bc* genes and captured an *idefix* and *17.6* retroviral insertion in antisense orientation (indicated insertion ages based on pairwise LTR divergence estimates). **(E)** Shown are microRNA-normalized levels of ovarian piRNAs from indicated wildtype strains (*iso1*, *w^1118^*) and cluster *77B* mutants (*77B^m1^*and *77B^minos^*). The transcription start site (TSS) of the cluster within the *oquasi* LTR is indicated (image in scale and aligned to panel D). **(F)** RNA-smFISH based expression analysis of the somatic *77B* piRNA cluster (black) in the germarium. GFP-trap lines indicate escort cells and early follicle cells, respectively (scale bar: 10µm). **(G)** Shown are X-Gal stainings of ovaries harboring lacZ-reporter transgenes for *flamenco* or cluster *77B* (arrow heads indicate specific signal in the germarium; scale bars: 100 μm). The detailed confocal analysis of the cluster *77B*-reporter in the germarium shows lacZ expression in black and DNA in magenta (Hoechst). **(H)** RNA-smFISH based expression analysis of the *17.6* retrovirus (black) in germaria (with indicated cell types labeled by GFP-trap lines) of *77B^m1-1^/w^1118^*control (top) ovaries or of *cluster 77B* mutant ovaries (trans-heterozygous for independent alleles *77B^m1^/^m2^*) (bottom; scale bar: 10µm).

Notably, *flamenco* harbors moderate silencing potential against the retroviruses *idefix* and *quasimodo* and none against *17.6* (Fig 5B) . Despite this, *17.6* was repressed in a piRNA pathway-dependent manner (Suppl. Fig. 11A), and wildtype ovaries contained abundant somatic antisense piRNAs against this retrovirus (Fig. 5C), suggesting the presence of an unidentified piRNA cluster responsible for *17.6* silencing. We examined *17.6* insertions in the reference genome and noticed an intriguing locus on chromosome 3L at cytological position 77B, that contains three retroviral insertions belonging to the *idefix* subclade (Fig. 5B, D) (*62*). This locus, referred to as *cluster 77B,* harbors a partial copy of an old *quasimodo*-related element (*oquasi*) containing LTR and 5’ UTR sequences, as well as one insertion each of *17.6* and *idefix* inserted in opposite orientation to *oquasi*. Small RNA-seq data revealed that *cluster 77B* is unidirectionally expressed in ovaries, generating piRNAs associated with somatic Piwi but not with germline Piwi/Aub/Ago3 (Fig. 5E; Suppl. Fig. 16A-E). We found *cluster 77B* to be also expressed in cultured ovarian somatic cells (OSCs), and RNA-seq, PRO-seq, and small RNA-seq data collectively indicate that its transcription start site corresponds to the initiator-motif within the first *oquasi* LTR, from where a polyadenylated transcript approximately 17 kb in length is generated (Suppl. Fig. 16F).

In contrast to the broad soma-expression of *flamenco*, *cluster 77B* exhibited exclusive expression in escort cells and early follicle cells within the germarium (Fig. 5F), closely resembling the expression of the *quasimodo* retrovirus (Suppl. Fig. 10). To extend these findings, we generated *flamenco* and *cluster 77B* reporter transgenes that incorporated lacZ under the control of the *flamenco* promoter region or the *oquasi* LTR plus 5’ UTR regions, respectively. In support of the smFISH data, the *flamenco* reporter was expressed in all somatic cells of the ovary, while the *cluster 77B* reporter recapitulated the specific expression pattern of the endogenous piRNA cluster in the germarium (Fig. 5G). Thus, *D. melanogaster* has co-opted the LTR and 5’ UTR sequences of the ancestral retrovirus *oquasi* to facilitate the transcription of its own silencing program, the evolutionarily young *cluster 77B*.

The expression domain of *cluster 77B* overlapped with that of the *17.6* retrovirus, for which *flamenco* lacks silencing potential. To investigate the functional significance of *cluster 77B*, we generated a mutant allele by deleting the first *oquasi* LTR, harboring the transcription start site. Flies homozygous for this allele, as well as those carrying a Minos insertion within the *oquasi* 5’ region, exhibited a loss of detectable *cluster 77B* expression and corresponding piRNA populations (Fig. 5E; Suppl. Fig. 16G). In these mutant flies, *17.6* was specifically de-repressed in escort cells and early follicle cells, highlighting the critical role of *cluster 77B* in silencing this retrovirus (Fig. 5H).

Finally, we investigated the evolutionary history of *cluster 77B*. By analyzing pairwise LTR divergences, we estimated that the *oquasi* insertion, which is closely related to the *Drosophila erecta* retrovirus *Nquasimodo* (*29*), dates back over 500,000 years ago (a likely underestimate due to a partial deletion of the *oquasi* 3’ LTR). The *idefix* and *17.6* insertions occurred approximately 100,000 and 40,000 years ago, respectively. Consistent with these timeframes, the syntenic genomic regions in *D. simulans* and *D. sechellia*, which share a common ancestor with *D. melanogaster* around 2.7 million years ago (*63*), contain the *oquasi* fragment but lack the *17.6* and *idefix* insertions (Fig. 5D). Since the *17.6* retrovirus is absent from the genomes of *D. simulans and D. sechellia*, *cluster 77B* appears to have evolved as a specific defense strategy against *17.6* in *D. melanogaster*. We therefore examined the resource of geographical *D. melanogaster* isolates (DSPR founder strains) (*64*) and discovered extensive structural variation at *cluster 77B*, indicating an ongoing adaptive process. Notably, *cluster 77B* consistently exhibits a preference for antisense insertions of retroviruses with varying insertion ages, but almost always belonging to the *idefix* subclade, precisely those that are co-expressed with *cluster 77B* in the germarium (Suppl. Fig. 17).

In conclusion, retroviruses and the somatic piRNA pathway display an intricate and interrelated co-evolution: the *flamenco* master locus has progressively incorporated sequence fragments from nearly all retroviruses, and the evolutionarily young *cluster 77B* ensures optimal repression of the *idefix* subclade in the germarium. The extensive co-evolution of the somatic piRNA pathway strongly suggests that unrestricted iERV expression in different somatic cell types is detrimental and that silencing of the diversified iERVs in the soma confers a selective advantage to the host.

## Discussion

In this study, we investigated an entire clade of endogenous retroviruses and the host’s defense mechanisms against them in a single model organism. By combining the systematic analysis of iERV sequences with their phylogenetic relationships, infectivity status, and functional expression patterns, our work provides unparalleled insights into the evolutionary diversification of endogenous retroviruses and their adaptations to an animal gonadal “ecosystem” and the co-evolving piRNA pathway.

Our key finding is that while iERVs collectively occupy every cell niche in the *Drosophila* ovary, infectious iERVs have evolved distinct expression patterns in the soma and derived, non-infectious iERVs converge in germline expression. Although stochastic events cannot be ruled out, we argue that this diversification in expression patterns reflects adaptive processes. In analogy to evolutionary theory, it would represent an intriguing example of niche partitioning among TEs, exhibiting characteristics analogous to an adaptive radiation. An adaptive radiation is the evolution of ecological diversity within a rapidly reproducing lineage (*65*). It is generally triggered by ecological opportunities such as the colonization of a new habitat, the extinction of antagonists, or the emergence of a key innovation that confers adaptive advantages. Intrigued by the conceptual similarities, we discuss iERV evolution in light of the criteria that characterize an adaptive radiation: *common ancestry*, *phenotype-environment correlation*, *trait utility*, and *rapid speciation* (*65*).

Our independent phylogenetic analyses, based on Pol, Gag, and Env alignments, strongly support the monophyletic nature of the iERV clade. These findings are consistent with the hypothesis that an LTR retroelement that acquired *env-F* de novo was the *common ancestor* of iERVs (*20, 21*). iERVs are the most diverse LTR retrotransposon clade in *D. melanogaster* and its close relatives (*29*). Together with the comparatively shorter branch lengths of iERVs in the phylogenetic tree compared to other LTR clades (Fig. 1A), this indicates a plausible scenario of comparatively *rapid speciation*.

During *Drosophila* oogenesis, primordial germ cells and their differentiating progeny closely interact with diverse somatic cell types that regulate germ cell differentiation while also providing protection and nourishment to the germline. Our findings reveal the remarkable adaptation of iERVs to this complex ecosystem, with the traits of infectivity and expression pattern strictly co-evolving, thereby showing a clear *phenotype-environment correlation*. Our smFISH analysis throughout oogenesis and in early embryos revealed that retroviruses and retroelements have evolved different replication strategies to target the host germline genome. Together with previous observations that demonstrate amplification of non-infectious as well as infectious iERVs in flies (*47, 66*), these findings argue that iERV replication strategies strictly correlate with their respective traits *cis*-regulatory elements and *env-F* status (*trait utility*).

The acquisition of *env-F* was undoubtedly a pivotal event in the radiation of iERVs. This key innovation can be considered as the adaptive breakthrough that provided the ecological opportunity for an ancestral retrovirus to colonize a new and diverse ecological niche, the ovarian soma. We propose that this diverse and initially unprotected habitat allowed infectious iERVs to readily replicate and diversify. This likely resulted in competition among iERVs for shard host resources and inter-viral interference during viral particle assembly (*67–70*). As a result, iERVs might have adapted their somatic expression patterns to discrete niches, possibly also adapting to different receptors expressed along the germline differentiation axis (Suppl. Fig. 18). Our data identify the co-evolving piRNA pathway as yet another force impacting iERV evolution. As somatic piRNA clusters with an insertion of an infectious iERV will shut down retroviral replication, such upgrades of host defense may have constituted a strong selection pressure, fostering the emergence of germline expressed, non-infectious retroelements. Loss of functional *env-F* occurred either passively as it was not anymore required for replication, or actively due to *env-F* expression being disadvantageous for germline replication or even detrimental for host germline development.

Three additional factors likely further promoted the adaptive radiation of iERVs. First, gain of infectivity significantly enlarged the replication habitat of iERVs. While the ovarian germline consists of only two transcriptionally distinct cell types (germline stem cells and differentiating nurse cells), the ovarian soma comprises at least eight cell types with diverse gene expression profiles (*71, 72*). This difference in cell-type diversity likely contributed to retroviruses diverging more readily compared to retroelement descendants. Secondly, the diversification rate of iERVs has probably been influenced by the adaptive capacity of the host defense systems. Lacking Rhino, the somatic piRNA pathway is less adaptive than its germline counterpart, giving retroviruses potentially more opportunities to diversify and adapt to their environment compared to retroelements. The third point pertains to the process of iERV diversification. Considering that the piRNA pathway effectively suppresses iERV expression, it can be inferred that iERV diversification must have occurred during periods of uncontrolled replication. Once a host population established piRNA-mediated control over a specific iERV lineage, further diversification could only take place if there was a temporary loss of piRNA control or through horizontal transfer of an iERV lineage to a naïve host population or species. In fact, over evolutionary timescales, horizontal transfer is a common phenomenon among TEs in *Drosophila*, including iERVs (*29*), indicating that iERVs diversified within a host range comprising multiple *Drosophila* species that share overlapping habitats.

Finally, our findings significant implications for the broader domain of LTR retrotransposon evolution. Firstly, niche partitioning, which indicates an adaptive evolutionary process driven by competition, is not exclusive to iERVs. For instance, in *Saccharomyces cerevisiae*, the LTR retrotransposons *Ty1* and *Ty3* have partitioned the single-celled host into discrete retro-transposition niches, with *Ty3* being expressed in haploid cells and *Ty1* in diploid cells (*73*). In vertebrates, infectious retroviruses of the *Retroviridae*, with a different origin compared to iERVs, have co-evolved with their hosts for millions of years. Due to stochastic germline infections, the *Retroviridae* have given rise to multiple lineages of endogenous retroviruses (ERVs) in vertebrate genomes (*48*). Notably, several vertebrate ERVs are expressed during specific developmental time windows or conditions. For example, during human preimplantation development, the LTRs of different HERV lineages exhibit activity in striking, often non-overlapping spatiotemporal patterns (*74, 75*). Secondly, several vertebrate ERVs also display a loss of their *env* gene. An example is the intracisternal A particle (IAP) in mice, a highly active ERV lineage, which evolved from a retroviral form, IAP-E, that encoded a functional *env* gene (*76*). Similar to non-infectious iERVs in *Drosophila*, IAPs are expressed in the mouse germline (*77*) and replicate successfully despite being targeted by the piRNA pathway (*78, 79*). Thus, the ecological and evolutionary principles underlying the diversification of iERVs in *Drosophila* likely also impact the evolution of vertebrate ERVs and their adaptations to specific host environments. In this context, our findings provide critical insights into the origins of eukaryotic *cis*-regulatory elements, which have repeatedly been co-opted from LTR regulatory sequences (*80*).

## Acknowledgements

We thank the VBCF core facilities (NGS, VDRC) for sequencing and fly stocks, the in-house Protein Chemistry and BioOptics facility for excellent support, and the Fly&Worm Facility for transgenesis and CRISPR-mediated genome engineering. We thank Thomas Harivel, Matthias Schäfer, and Maria Novatchkova for experimental and analytical support, and Chantal Vaury, Jereoen Dobblare, the Bloomington and VDRC stock centers for flies and the Developmental Studies Hybridoma Bank (DSHB) for antibodies. We thank members of the Institute of Population Genetics (VetMed University Vienna), the Brennecke laboratory, and Andrea Betancourt for discussions, and Life Science Editors, Andrea Pauli, Ortrun Mittelsten Scheid, Alejandro Burga, Arturo Marí-Ordóñez, Alexander Hayward, and Justin Blumenstiel for comments on the manuscript.

## Author contributions

KAS: conception, experiments, analysis, writing, supervision, funding; DH: small-RNA-libraries, computational analysis of NGS and genomics data; BR: experiments; CK: phylogenetic analysis; CS: funding & infrastructure; JB: analysis, writing, supervision, funding & infrastructure.

## Data and Materials availability

All fly stocks generated for this study are available from VDRC, antibodies are available upon request, NGS data is available from GEO.

## Supplementary Materials

Materials and Methods

Figs. S1 to S18

Table S1: fly stocks, primers, FISH probes, antibodies, NGS libraries, & piRNA cluster coordinates

Document S1: lists all iERV consensus sequences used in this study

## References

1. G. Bourque et al., Ten things you should know about transposable elements. Genome Biol 19, 199 (2018).

2. C. D. Malone, G. J. Hannon, Small RNAs as guardians of the genome. Cell 136, 656–668 (2009).

3. H. L. Levin, J. V. Moran, Dynamic interactions between transposable elements and their hosts. Nat Rev Genet 12, 615–627 (2011).

4. N. V. Fedoroff, Presidential address. Transposable elements, epigenetics, and genome evolution. Science 338, 758–767 (2012).

5. M. G. Kidwell, D. R. Lisch, Perspective: transposable elements, parasitic DNA, and genome evolution. Evolution 55, 1–24 (2001).

6. M. G. Kidwell, Transposable elements and the evolution of genome size in eukaryotes. Genetica 115, 49–63 (2002).

7. A. Bohne, F. Brunet, D. Galiana-Arnoux, C. Schultheis, J. N. Volff, Transposable elements as drivers of genomic and biological diversity in vertebrates. Chromosome Res 16, 203–215 (2008).

8. G. Bourque, Transposable elements in gene regulation and in the evolution of vertebrate genomes. Curr Opin Genet Dev 19, 607–612 (2009).

9. N. J. Bowen, I. K. Jordan, Transposable elements and the evolution of eukaryotic complexity. Curr Issues Mol Biol 4, 65–76 (2002).

10. C. Feschotte, E. J. Pritham, DNA transposons and the evolution of eukaryotic genomes. Annu Rev Genet 41, 331–368 (2007).

11. R. L. Cosby, N. C. Chang, C. Feschotte, Host-transposon interactions: conflict, cooperation, and cooption. Genes Dev 33, 1098–1116 (2019).

12. H. D. Madhani, The frustrated gene: origins of eukaryotic gene expression. Cell 155, 744–749 (2013).

13. C. Feschotte, Transposable elements and the evolution of regulatory networks. Nat Rev Genet 9, 397–405 (2008).

14. S. D. Fugmann, A. I. Lee, P. E. Shockett, I. J. Villey, D. G. Schatz, The RAG proteins and V(D)J recombination: complexes, ends, and transposition. Annu Rev Immunol 18, 495–527 (2000).

15. S. Mi et al., Syncytin is a captive retroviral envelope protein involved in human placental morphogenesis. Nature 403, 785–789 (2000).

16. C. Llorens, B. Soriano, M. Krupovic, C. Ictv Report, ICTV Virus Taxonomy Profile: Metaviridae. J Gen Virol 101, 1131–1132 (2020).

17. Y. Stefanov, V. Salenko, I. Glukhov, Drosophila errantiviruses. Mob Genet Elements 2, 36–45 (2012).

18. K. A. Senti, J. Brennecke, The piRNA pathway: a fly’s perspective on the guardian of the genome. Trends Genet 26, 499–509 (2010).

19. M. C. Siomi, K. Sato, D. Pezic, A. A. Aravin, PIWI-interacting small RNAs: the vanguard of genome defence. Nat Rev Mol Cell Biol 12, 246–258 (2011).

20. C. Terzian, A. Pelisson, A. Bucheton, Evolution and phylogeny of insect endogenous retroviruses. BMC Evol Biol 1, 3 (2001).

21. H. S. Malik, S. Henikoff, T. H. Eickbush, Poised for contagion: evolutionary origins of the infectious abilities of invertebrate retroviruses. Genome Res 10, 1307–1318 (2000).

22. G. F. Rohrmann, P. A. Karplus, Relatedness of baculovirus and gypsy retrotransposon envelope proteins. BMC Evol Biol 1, 1 (2001).

23. A. Pelisson et al., Gypsy transposition correlates with the production of a retroviral envelope-like protein under the tissue-specific control of the Drosophila flamenco gene. Embo J 13, 4401–4411 (1994).

24. S. U. Song, T. Gerasimova, M. Kurkulos, J. D. Boeke, V. G. Corces, An env-like protein encoded by a Drosophila retroelement: evidence that gypsy is an infectious retrovirus. Genes Dev 8, 2046–2057 (1994).

25. P. Leblanc et al., Life cycle of an endogenous retrovirus, ZAM, in Drosophila melanogaster. J Virol 74, 10658–10669 (2000).

26. E. Brasset et al., Viral particles of the endogenous retrovirus ZAM from Drosophila melanogaster use a pre-existing endosome/exosome pathway for transfer to the oocyte. Retrovirology 3, 25 (2006).

27. V. V. Kapitonov, J. Jurka, Molecular paleontology of transposable elements in the Drosophila melanogaster genome. Proc Natl Acad Sci U S A 100, 6569–6574 (2003).

28. J. S. Kaminker et al., The transposable elements of the Drosophila melanogaster euchromatin: a genomics perspective. Genome Biol 3, RESEARCH0084 (2002).

29. N. Bargues, E. Lerat, Evolutionary history of LTR-retrotransposons among 20 Drosophila species. Mob DNA 8, 7 (2017).

30. S. Tcheressiz et al., Expression of the Idefix retrotransposon in early follicle cells in the germarium of Drosophila melanogaster is determined by its LTR sequences and a specific genomic context. Mol Genet Genomics 267, 133–141 (2002).

31. R. M. Marsano et al., The complete Tirant transposable element in Drosophila melanogaster shows a structural relationship with retrovirus-like retrotransposons. Gene 247, 87–95 (2000).

32. S. Shigenobu, Y. Kitadate, C. Noda, S. Kobayashi, Molecular characterization of embryonic gonads by gene expression profiling in Drosophila melanogaster. Proc Natl Acad Sci U S A 103, 13728–13733 (2006).

33. M. Westenberg et al., Furin is involved in baculovirus envelope fusion protein activation. J Virol 76, 178–184 (2002).

34. M. Westenberg et al., Functional analysis of the putative fusion domain of the baculovirus envelope fusion protein F. J Virol 78, 6946–6954 (2004).

35. R. G. FitzJohn, W. P. Maddison, S. P. Otto, Estimating trait-dependent speciation and extinction rates from incompletely resolved phylogenies. Syst Biol 58, 595–611 (2009).

36. M. P. Davis, P. E. Midford, W. Maddison, Exploring power and parameter estimation of the BiSSE method for analyzing species diversification. BMC Evol Biol 13, 38 (2013).

37. D. Kirilly, T. Xie, The Drosophila ovary: an active stem cell community. Cell Res 17, 15–25 (2007).

38. J. C. Duhart, T. T. Parsons, L. A. Raftery, The repertoire of epithelial morphogenesis on display: Progressive elaboration of Drosophila egg structure. Mech Dev 148, 18–39 (2017).

39. C. D. Malone et al., Specialized piRNA pathways act in germline and somatic tissues of the Drosophila ovary. Cell 137, 522–535 (2009).

40. G. Dietzl et al., A genome-wide transgenic RNAi library for conditional gene inactivation in Drosophila. Nature 448, 151–156 (2007).

41. J. Q. Ni et al., A genome-scale shRNA resource for transgenic RNAi in Drosophila. Nat Methods 8, 405–407 (2011).

42. D. Handler et al., The Genetic Makeup of the Drosophila piRNA Pathway. Mol Cell 50, 762–777 (2013).

43. K. A. Senti, D. Jurczak, R. Sachidanandam, J. Brennecke, piRNA-guided slicing of transposon transcripts enforces their transcriptional silencing via specifying the nuclear piRNA repertoire. Genes Dev 29, 1747–1762 (2015).

44. A. Raj, P. van den Bogaard, S. A. Rifkin, A. van Oudenaarden, S. Tyagi, Imaging individual mRNA molecules using multiple singly labeled probes. Nat Methods 5, 877–879 (2008).

45. A. S. Raikhel, T. S. Dhadialla, Accumulation of yolk proteins in insect oocytes. Annu Rev Entomol 37, 217–251 (1992).

46. Y. Hara, D. Yamamoto, Effects of Food and Temperature on Drosophila melanogaster Reproductive Dormancy as Revealed by Quantification of a GFP-Tagged Yolk Protein in the Ovary. Front Physiol 12, 803144 (2021).

47. L. Wang, K. Dou, S. Moon, F. J. Tan, Z. Z. Zhang, Hijacking Oogenesis Enables Massive Propagation of LINE and Retroviral Transposons. Cell 174, 1082–1094 e1012 (2018).

48. W. E. Johnson, Origins and evolutionary consequences of ancient endogenous retroviruses. Nat Rev Microbiol 17, 355–370 (2019).

49. S. Desset, C. Meignin, B. Dastugue, C. Vaury, COM, a heterochromatic locus governing the control of independent endogenous retroviruses from Drosophila melanogaster. Genetics 164, 501–509 (2003).

50. C. Klattenhoff et al., The Drosophila HP1 homolog Rhino is required for transposon silencing and piRNA production by dual-strand clusters. Cell 138, 1137–1149 (2009).

51. Z. Zhang et al., The HP1 homolog rhino anchors a nuclear complex that suppresses piRNA precursor splicing. Cell 157, 1353–1363 (2014).

52. F. Mohn, G. Sienski, D. Handler, J. Brennecke, The rhino-deadlock-cutoff complex licenses noncanonical transcription of dual-strand piRNA clusters in Drosophila. Cell 157, 1364–1379 (2014).

53. D. Gebert et al., Large Drosophila germline piRNA clusters are evolutionarily labile and dispensable for transposon regulation. Mol Cell 81, 3965–3978 e3965 (2021).

54. I. Olovnikov et al., De novo piRNA cluster formation in the Drosophila germ line triggered by transgenes containing a transcribed transposon fragment. Nucleic Acids Res 41, 5757–5768 (2013).

55. S. Shpiz, S. Ryazansky, I. Olovnikov, Y. Abramov, A. Kalmykova, Euchromatic Transposon Insertions Trigger Production of Novel Pi- and Endo-siRNAs at the Target Sites in the Drosophila Germline. PLoS Genet 10, e1004138 (2014).

56. L. Baumgartner et al., The Drosophila ZAD zinc finger protein Kipferl guides Rhino to piRNA clusters. Elife 11, (2022).

57. C. Goriaux, S. Desset, Y. Renaud, C. Vaury, E. Brasset, Transcriptional properties and splicing of the flamenco piRNA cluster. EMBO Rep 15, 411–418 (2014).

58. E. Sarot, G. Payen-Groschene, A. Bucheton, A. Pelisson, Evidence for a piwi-dependent RNA silencing of the gypsy endogenous retrovirus by the Drosophila melanogaster flamenco gene. Genetics 166, 1313–1321 (2004).

59. S. Desset et al., Mobilization of two retroelements, ZAM and Idefix, in a novel unstable line of Drosophila melanogaster. Mol Biol Evol 16, 54–66 (1999).

60. J. Brennecke et al., Discrete Small RNA-Generating Loci as Master Regulators of Transposon Activity in Drosophila. Cell 128, 1089–1103 (2007).

61. V. Zanni et al., Distribution, evolution, and diversity of retrotransposons at the flamenco locus reflect the regulatory properties of piRNA clusters. Proc Natl Acad Sci U S A 110, 19842–19847 (2013).

62. P. Chen, A. A. Aravin, Genetic control of a sex-specific piRNA program. Curr Biol 33, 1825–1835 e1823 (2023).

63. D. J. Obbard et al., Estimating divergence dates and substitution rates in the Drosophila phylogeny. Mol Biol Evol 29, 3459–3473 (2012).

64. M. Chakraborty, J. J. Emerson, S. J. Macdonald, A. D. Long, Structural variants exhibit widespread allelic heterogeneity and shape variation in complex traits. Nat Commun 10, 4872 (2019).

65. D. Schluter, The Ecology of Adaptive Radiation. Oxford Series in Ecology & Evolution (2000).

66. B. Barckmann et al., The somatic piRNA pathway controls germline transposition over generations. Nucleic Acids Res 46, 9524–9536 (2018).

67. M. Mura et al., Late viral interference induced by transdominant Gag of an endogenous retrovirus. Proc Natl Acad Sci U S A 101, 11117–11122 (2004).

68. D. Trono, M. B. Feinberg, D. Baltimore, HIV-1 Gag mutants can dominantly interfere with the replication of the wild-type virus. Cell 59, 113–120 (1989).

69. M. A. Sommerfelt, R. A. Weiss, Receptor interference groups of 20 retroviruses plating on human cells. Virology 176, 58–69 (1990).

70. M. C. Rosales Gerpe et al., The U3 and Env Proteins of Jaagsiekte Sheep Retrovirus and Enzootic Nasal Tumor Virus Both Contribute to Tissue Tropism. Viruses 11, (2019).

71. T. D. Hinnant, J. A. Merkle, E. T. Ables, Coordinating Proliferation, Polarity, and Cell Fate in the Drosophila Female Germline. Front Cell Dev Biol 8, 19 (2020).

72. M. Slaidina, S. Gupta, T. U. Banisch, R. Lehmann, A single-cell atlas reveals unanticipated cell type complexity in Drosophila ovaries. Genome Res 31, 1938–1951 (2021).

73. S. Sandmeyer, K. Patterson, V. Bilanchone, Ty3, a Position-specific Retrotransposon in Budding Yeast. Microbiol Spectr 3, MDNA3-0057-2014 (2015).

74. J. Goke et al., Dynamic transcription of distinct classes of endogenous retroviral elements marks specific populations of early human embryonic cells. Cell Stem Cell 16, 135–141 (2015).

75. T. A. Carter et al., Mosaic cis-regulatory evolution drives transcriptional partitioning of HERVH endogenous retrovirus in the human embryo. Elife 11, (2022).

76. D. Ribet et al., An infectious progenitor for the murine IAP retrotransposon: emergence of an intracellular genetic parasite from an ancient retrovirus. Genome Res 18, 597–609 (2008).

77. A. Dupressoir, T. Heidmann, Germ line-specific expression of intracisternal A-particle retrotransposons in transgenic mice. Mol Cell Biol 16, 4495–4503 (1996).

78. M. A. Carmell et al., Miwi2 is essential for spermatogenesis and repression of transposons in the mouse male germline. Dev Cell in press, (2007).

79. G. Magiorkinis, R. J. Gifford, A. Katzourakis, J. De Ranter, R. Belshaw, Env-less endogenous retroviruses are genomic superspreaders. Proc Natl Acad Sci U S A 109, 7385–7390 (2012).

80. E. B. Chuong, N. C. Elde, C. Feschotte, Regulatory activities of transposable elements: from conflicts to benefits. Nat Rev Genet 18, 71–86 (2017).

81. W. Bao, K. K. Kojima, O. Kohany, Repbase Update, a database of repetitive elements in eukaryotic genomes. Mob DNA 6, 11 (2015).

82. V. V. Kapitonov, J. Jurka, A universal classification of eukaryotic transposable elements implemented in Repbase. Nat Rev Genet 9, 411–412; author reply 414 (2008).

83. V. Ranwez, E. J. P. Douzery, C. Cambon, N. Chantret, F. Delsuc, MACSE v2: Toolkit for the Alignment of Coding Sequences Accounting for Frameshifts and Stop Codons. Mol Biol Evol 35, 2582–2584 (2018).

84. J. J. Almagro Armenteros et al., SignalP 5.0 improves signal peptide predictions using deep neural networks. Nat Biotechnol 37, 420–423 (2019).

85. J. J. Almagro Armenteros et al., Detecting sequence signals in targeting peptides using deep learning. Life Sci Alliance 2, (2019).

86. A. Krogh, B. Larsson, G. von Heijne, E. L. Sonnhammer, Predicting transmembrane protein topology with a hidden Markov model: application to complete genomes. J Mol Biol 305, 567–580 (2001).

87. A. Stamatakis, RAxML version 8: a tool for phylogenetic analysis and post-analysis of large phylogenies. Bioinformatics 30, 1312–1313 (2014).

88. J. Trifinopoulos, L. T. Nguyen, A. von Haeseler, B. Q. Minh, W-IQ-TREE: a fast online phylogenetic tool for maximum likelihood analysis. Nucleic Acids Res 44, W232–235 (2016).

89. S. Q. Le, O. Gascuel, An improved general amino acid replacement matrix. Mol Biol Evol 25, 1307–1320 (2008).

90. D. H. Huson, C. Scornavacca, Dendroscope 3: an interactive tool for rooted phylogenetic trees and networks. Syst Biol 61, 1061–1067 (2012).

91. R. C. Edgar, MUSCLE: multiple sequence alignment with high accuracy and high throughput. Nucleic Acids Res 32, 1792–1797 (2004).

92. N. J. Bowen, J. F. McDonald, Drosophila euchromatic LTR retrotransposons are much younger than the host species in which they reside. Genome Res 11, 1527–1540 (2001).

93. P. D. Keightley et al., Analysis of the genome sequences of three Drosophila melanogaster spontaneous mutation accumulation lines. Genome Res 19, 1195–1201 (2009).

94. A. D. Cutter, Divergence times in Caenorhabditis and Drosophila inferred from direct estimates of the neutral mutation rate. Mol Biol Evol 25, 778–786 (2008).

95. F. Sievers et al., Fast, scalable generation of high-quality protein multiple sequence alignments using Clustal Omega. Mol Syst Biol 7, 539 (2011).

96. J. Dobbelaere, T. Y. Su, B. Erdi, A. Schleiffer, A. Dammermann, A phylogenetic profiling approach identifies novel ciliogenesis genes in Drosophila and C. elegans. EMBO J, e113616 (2023).

97. X. Morin, R. Daneman, M. Zavortink, W. Chia, A protein trap strategy to detect GFP-tagged proteins expressed from their endogenous loci in Drosophila. Proc Natl Acad Sci U S A 98, 15050–15055 (2001).

98. M. Buszczak et al., The carnegie protein trap library: a versatile tool for Drosophila developmental studies. Genetics 175, 1505–1531 (2007).

99. Z. Zhang, W. E. Theurkauf, Z. Weng, P. D. Zamore, Strand-specific libraries for high throughput RNA sequencing (RNA-Seq) prepared without poly(A) selection. Silence 3, 9 (2012).

100. I. Gaspar, F. Wippich, A. Ephrussi, Enzymatic production of single-molecule FISH and RNA capture probes. RNA 23, 1582–1591 (2017).

101. G. L. Glotzer, P. Tardivo, E. M. Tanaka, Canonical Wnt signaling and the regulation of divergent mesenchymal Fgf8 expression in axolotl limb development and regeneration. Elife 11, (2022).

102. T. Trcek, T. Lionnet, H. Shroff, R. Lehmann, mRNA quantification using single-molecule FISH in Drosophila embryos. Nat Protoc 12, 1326–1348 (2017).

103. T. Doura et al., Detection of LacZ-Positive Cells in Living Tissue with Single-Cell Resolution. Angew Chem Int Ed Engl 55, 9620–9624 (2016).

104. R. A. Hoskins et al., A BAC-based physical map of the major autosomes of Drosophila melanogaster. Science 287, 2271–2274 (2000).

105. K. J. Venken et al., Versatile P[acman] BAC libraries for transgenesis studies in Drosophila melanogaster. Nat Methods 6, 431–434 (2009).

106. M. Markstein, C. Pitsouli, C. Villalta, S. E. Celniker, N. Perrimon, Exploiting position effects and the gypsy retrovirus insulator to engineer precisely expressed transgenes. Nat Genet 40, 476–483 (2008).

107. D. T. Ge, C. Tipping, M. H. Brodsky, P. D. Zamore, Rapid Screening for CRISPR-Directed Editing of the Drosophila Genome Using white Coconversion. G3 (Bethesda) 6, 3197–3206 (2016).

108. T. Grentzinger et al., A universal method for the rapid isolation of all known classes of functional silencing small RNAs. Nucleic Acids Res 48, e79 (2020).

109. C.. Drosophila 12 Genomes et al., Evolution of genes and genomes on the Drosophila phylogeny. Nature 450, 203–218 (2007).

110. M. Chakraborty et al., Hidden genetic variation shapes the structure of functional elements in Drosophila. Nat Genet 50, 20–25 (2018).

