## Supplementary Information for "Functional Adaptations of Endogenous Retroviruses to the *Drosophila* Host Underlie their Evolutionary Diversification"

**This PDF file includes:**
Materials and Methods
Figs. S1 to S18

#### MATERIALS AND METHODS

##### Annotation of LTR retrotransposon consensus sequences

Consensus sequences for all *Metaviridae* and *Belpaoviridae* LTR retrotransposons in *Drosophila melanogaster* were collected from the BDGP and Repbase databases as well as from primary literature (*28, 43, 81, 82*). In most cases, existing annotations for LTR, 5' UTR, and coding sequences of *gag*, *pol,* and *env-F* were adopted. For several retrotransposons, we manually curated the consensus sequences. For this, existing consensus sequences were used to search the dm6 reference genome for the longest and most complete insertions of the respective lineage. Within these, we identified sequence stretches representing open reading frames for *gag*, *pol*, and *env-F*, and aligned those with MASCE (*83*) to all other *gypsy/gypsy* clade sequences for the same ORF. This allowed the correction of isolated SNPs or small INDELs in *env-F* for the following consensus sequences: *gtwin*, *gypsy5*, *rover*, *HMS Beagle2*. For alignments containing generally incomplete ORFs (inactive lineages), additional shorter fragments were identified in dm6 and used to conservatively correct and fill gaps, frame shifts and stop codons, always using MACSE alignments. This process was reiterated until the respective consensus sequence could not be improved further using additional fragments. The curated ORF consensus sequences were used to assemble full length consensus sequences that were used in this study. The following consensus sequences were curated for multiple ORFs: *accord2*, *gypsy2*, *gypsy3*, *gypsy7*, *gypsy9*, *gypsy10*. To annotate the complete *env-F* coding sequences, splice junctions of sub-genomic *env-F* transcripts were determined. Total RNA was extracted from ovaries of *traffic jam*-GAL4 UAS-*vreteno*$^{GD}$ females (fly stocks listed in Table S1), PCR products were amplified using random primed first strand cDNA and primers flanking known and anticipated *env-F* splice junctions (primer sequences listed in Table S1). PCR amplicons were subcloned and sequenced. From the resulting full-length Env-F protein sequences, signal peptide and transmembrane domains were predicted with signalP 5.0 and target 1.1 or TMHMM 2.0, respectively (*84-86*). All consensus sequences used in the study are provided in Document S1. Based on the curated set of consensus sequences, we determined which lineages were represented by at least one structurally intact (active lineage) or exclusively structurally defective copies (inactive lineage) in the dm6 reference genome. For all retro-element consensus sequences lacking full-length *env-F*, the sequence from the end of *pol* extending into the 3′ LTR were aligned to full-length *env-F* sequences from the same subclade using MASCE to identify *env-F* ORF fragments.

**Sequence alignments and phylogenetic analyses**

The open reading frames of *gag*, *pol* and *env-F* in the curated consensus sequences were aligned using MACSE2.0 (*83*). Sequences for *gag* were restricted to the central core domain, sequences for *pol* to the Protease domain until the end of the conserved Integrase domain, sequences for *env-F* were trimmed at the N-terminus until the beginning of the signal peptide (*idefix* subclade). The resulting MACSE alignments were used to estimate phylogenetic trees using three methods, RAxML and IQ-tree (*87, 88*). For RAxML, the best-scoring substitution model was determined among the amino acid models as LG+F+Γ (LG with empirical base frequencies and the Γ model of rate heterogeneity; (*89*)), and clade support values were calculated from non-parametric bootstrapping implemented in RAxML based on 1000 replicates. The phylogenetic relationships were confirmed by performing phylogenetic analysis with and without the outgroups in IQ-TREE with 1000 ultrafast bootstraps. Phylogenetic trees were visualized and analyzed using Dendroscope (*90*). Diversitree (*35*) was used for the statistical analysis of the ancestral character state using the RAxML Pol tree without the *gypsy*/*mdg3* outgroup and states of either functional *env-F* or clear *env-F* fragments versus those lacking clear env-F evidence (*gypsy7* and *transpac*) at the terminal branches. BiSSE was used to estimate if speciation and extinction rates of retroviruses (with functional *env-F*) and retro-elements (without functional *env-F*) differed. LTR alignments were performed using MUSCLE alignments in MegAlign (DNASTAR) or the web interface at www.ebi.ac.at (*91*). The approximate ages of individual LTR-retrotransposon insertions in dm6 by pairwise LTR comparisons were estimated using the formula T=K/2xr (*92*) with r=0.0346, assuming ten generations per year (*63, 93, 94*). The nucleotide sequence divergence of individual iERVs within their respective subclades was assessed by extracting LTR, 5′ UTR, and ORFs for *gag* (core), *pol* (RT-RNAseH domain), and *env-F* from the respective consensus sequences, aligning them with MUSCLE (for LTR) or MASCE (for ORFs), and determining the percent sequence identities and the percentage of aligned nucleotides in Clustal Omega (*95*). To detect remnant *env-F* ORFs in retroelements, the sequence interval from the 3′ end of *pol* and to the and including downstream LTR was extracted and aligned to the *env-F* ORFs of retroviruses in the same subclade using MACSE2.0.

**Genetic analyses**

All fly stocks used in this study are listed in Table S1. Tissue specific knockdowns of piRNA pathway genes in the ovarian soma or germline were performed as described previously (*40-43*). For knockdown experiments in the male germline (testes), we used a driver line combining NGT40 (2[nd] chromosome) and *bam*-GAL4:VP16 (3[rd] chromosome) (*96*) with shRNA-lines targeting *white* (control) or *aub*+*ago3*. To identify somatic cell types, GFP-trap lines (*97, 98*) for *CG14207* (*CB02069*), *Drat* (*CB03410*), *fax* (*CC01359*) and *fas3* (*G00258*), labelling terminal filament cells, cap cells, escort cells or germarium early follicle cells and polar cells, respectively, were combined with the *tj*-Gal4 driven knockdown system. For germline knockdowns, a *GFP-nup107* transgene was introduced for labeling the nuclear envelope (*43*).

**polyA RNA-seq**

Total RNA was extracted with TRIzol from ovaries of adult flies or from dechorionated and washed 0-1h old embryos (prior to the onset of zygotic transcription). After RNAeasy column purification with on-column DNAse I digest, eluted RNA was 2x polyA selected using magnetic oligo-dT beads and further processed as strand-specific RNA-seq libraries (*99*). All sequenced libraries are listed in Table S1.

**smFISH probes**

smFISH probes were designed as previously described (*43*) and ordered either directly from LGC/Stellaris or as DNA oligos that were subsequently labeled in-house (*100*). All probe sets are listed in Table S1. lacZ HCR probes were designed as in (*101*).

**Antibodies**

All antibodies used in this study are listed in Table S1. For the newly generated transposon antibodies, we selected antigens that show a high level of sequence difference within an iERV subclade (*McClintock* Gag: ITEAQTAENFRPQASEQANS; *gypsy* Gag: EAPKQKDPKEEYEKTAKAA). Peptides were synthesized with terminal cysteine residues and coupled to KLH. Polyclonal antisera were raised in rabbits using the rapid protocol (Eurogentec).

**smFISH, HCR, immuno-fluorescence, and ß-Gal stainings**

Ovary and testes single molecule fluorescent RNA in Situ Hybridization (smFISH) and antibody stainings in ovaries or embryos were performed as previously described (*43*). For smFISH/antibody co-stainings, we first performed RNA smFISH, followed by anti-Armadillo stainings in 2xSSC based buffers. RNA smFISH stainings in early embryos were according to (*102*). HCR was performed as in (*101*). Reporter transgene expressions were analyzed by *lacZ* RNA smFISH, *lacZ* HCR-FISH, or anti-β-Gal stainings. Chromogenic β-Galactosidase assays were performed with blue precipitate-forming X-Gal (*42*) or fluorescent SPIDER β-Gal substrate: Ovaries of lacZ-reporter expressing flies were fixed with 4% PFA in PBS at room temperature and stained in PBS containing 1ug/ml Hoechst 33342 and 2μM fluorescent SPiDER β-gal reagent at 37° Celsius for 30 minutes, followed by 3x PBS washes and mounting in Prolong Diamond mounting medium (*103*). All specimens were imaged on a Zeiss LSM780 confocal microscope or on an Axio Imager.Z2 widefield microscope with an Axiocam 506 color camera for X-gal stainings.

**LacZ reporter transgenes and functional analysis**

Transposon reporters were based on a vector backbone with attB, mini-white marker, multi cloning site, and a 3xHA-lacZ reporter followed by a 3´ UTR containing a short *ftz*-intron, and *p10* terminator. *cis*-regulatory sequences of individual iERVs were amplified from genomic BACR clones (*104*) or Pacman clones (*105*) containing a corresponding full-length iERV insertion. PCR amplicons included the entire 5´ LTR and the 5´ UTR plus the *gag* ATG. Sequence-verified reporter constructs were integrated into *attP40* ($2^{nd}$ chromosome) (*106*). Obtained reporter transgenes were combined with UAS transgenes for control or piRNA targeting RNAi constructs and crossed to the different Gal4 driver lines. The *flamenco* and *cluster 77B* transcriptional reporter constructs were generated similarly (using primers listed in Table S1) and were integrated into *attP2* (*flamenco*) and *attP40* (*cluster 77B*).

**piRNA cluster silencing potential**

Genomic sequences (dm6) from of the uni-strand piRNA clusters *flamenco* and *cluster 77B* and from the dual-strand piRNA clusters *38C, 42AB,* and *80F* were parsed into all possible 25mer sequences (mimicking piRNAs). These 25mer sequences were mapped to all iERV consensus sequences with no mismatches (similar results were obtained with three mismatches). The mappings along each consensus sequence were recorded in a binary fashion, resulting in the percentage of each iERV sequence that was complementary to cluster 25mers. Chromosomal coordinates (dm6) for the analyzed piRNA clusters are listed in Table S1.

##### *cluster 77B* mutagenesis

The presumptive promoter element of *cluster 77B* was deleted using CRISPR/HDR with two guide RNAs (GCTATGTAACGCCGTCTGCA and GCACAGCAAAATTTGCACTG) that were cloned into pCFD4D (*107*). The HDR repair template consisted of homology arms (784 bp left arm and 1022 bp right arm) flanking a GMR-*white*$^{+}$ marker that itself was flanked by FRT sites. Both plasmids were injected into embryos laid by *y,w*; *actin-Cas9* flies. Mutant alleles were sequence verified. The *FRT-GMR-white*$^{+}$*-FRT* cassette for *77B*$^{m1-1}$ was removed by crossing to a hs-Flp; Dr/TM3 strain and heat shocking. The Minos allele inside *cluster 77B* is *Mi(MIC)MI07908* and was obtained from the Bloomington stock center. All mutant alleles were crossed out for at least two generations to virgins of the *w; Dr/TM3,Sb* strain to homogenize their genetic background prior to phenotypic analysis.

##### Small RNA-seq libraries

Small RNA-seq libraries from ovaries were generated as previously described (*56*), taking advantage of the Argonaute isolation protocol via TraPR (*108*). All libraries are listed in Table S1.

##### Small RNA-seq analysis

Small-RNA-seq and PIWI protein IP small RNA-seq libraries were analyzed as previously (*43*), but using the updated iERV consensus sequences (no mismatches for mappings). Total small RNA-seq libraries analyzed were from ovaries of *MTD>rhino-sh* versus *MTD>white-sh* (control) (*52*) and from double mutants for the piRNA clusters *38C 42AB* versus *w*$^{1118}$ (control) (*53*). The piRNA analysis for *cluster 77B* mutants used genome-unique piRNAs mapping to piRNA *cluster 77B* with no mismatches. Tissue specific PIWI protein-IP-small RNA-seq libraries were analyzed as in (*43*).

##### Comparative analysis of piRNA *cluster 77B* in other species and *D. melanogaster* isolates

We analyzed the genomic organization of *cluster 77B* in *D. melanogaster* iso-1 (dm6), *D. simulans* and *D. sechellia* (*109*). For the strain analysis, we used assembled genomes of the starting strains of the *Drosophila* synthetic population resource (DSPR) (*110*) and annotated the content of the *cluster 77B* locus using a repeat masker analysis (*81*). The insertion ages of contained LTR-retrotransposons were calculated as described above.

Supplementary Figure 1

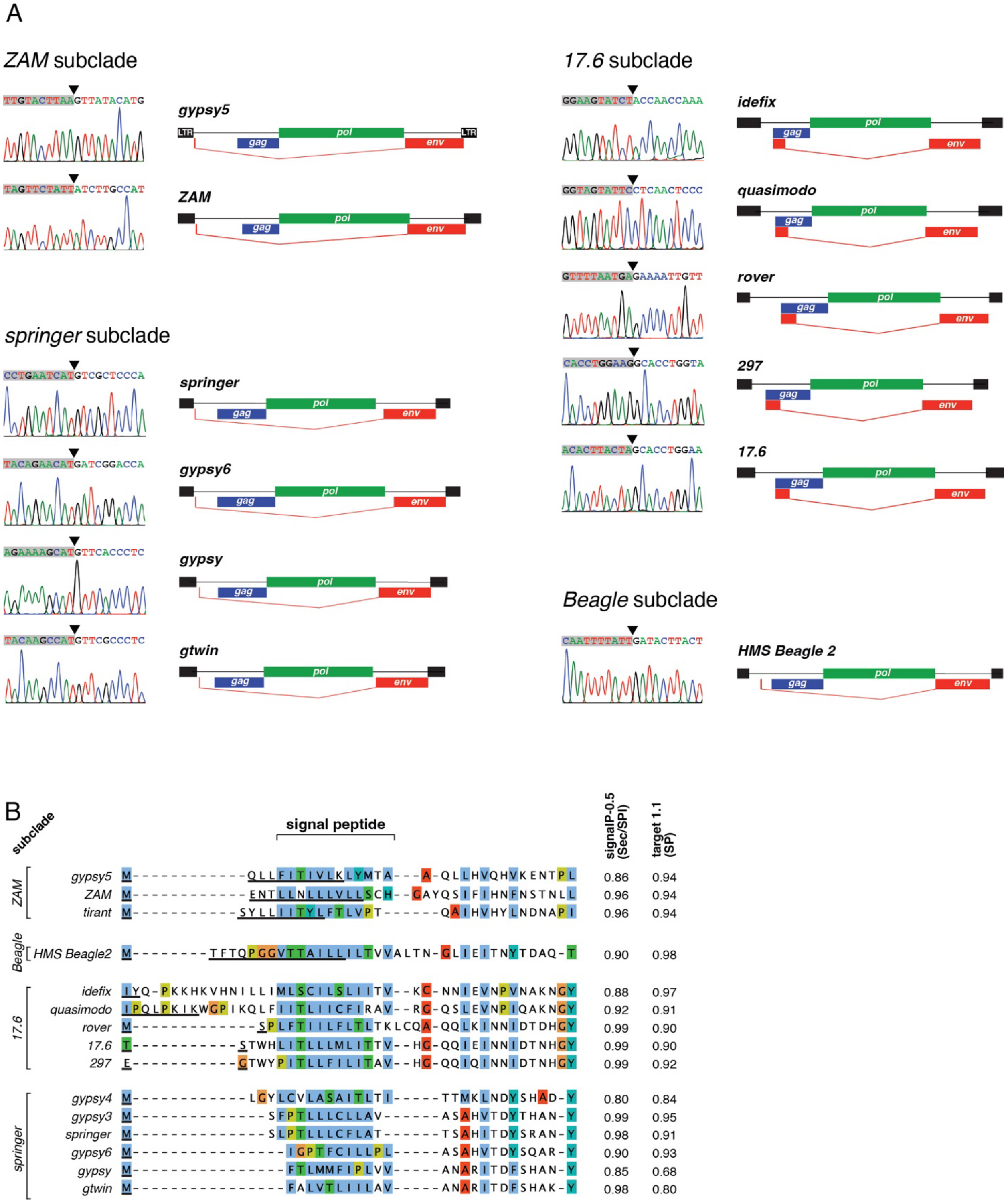

**Fig. S1: *env-F* splice patterns among infectious iERVs. (A)** Shown are Sanger sequencing tracks, centered on the *env-F* splice junction of indicated iERVs, from subcloned PCR products obtained from first strand cDNA prepared from total RNA of ovaries lacking somatic piRNA control (*tj*-Gal4 driven $vreteno^{GD}$ RNAi). Also shown are cartoons of the corresponding retrovirus indicating the *env-F* splice pattern, drawn to scale. **(B)** Protein sequence alignment of the spliced N-terminus of Env-F proteins from indicated iERV consensus sequences. Underlined amino acids are encoded by the upstream exon, signal peptide is indicated, predicted cleavage site (amino acid underlaid with red) and the respective scores using two different prediction algorithms are shown.

#### Supplementary Figure 2

**Functional retroviral envelope splice junctions and their conservation patterns**

| **ZAM subclade** | upstream | downstream | |
|---|---|---|---|
| ZAM (known, this study) | TAGTTCTATTGTAAGTAGTT | TGCATTCCAGATCTTGCCAT | seq confirmed |
| gypsy5 (this study) | TTGTACTTAAGTGAGTAGAG | ATCATTTCAGGTTATACATG | seq confirmed (distinct from consensus) |
| tirant (known) | TTACCTACCTGTAAGTAAAC | ACCATTTCAGGTTCACCCTT | not deregulated |

| **Idefix subclade** | upstream | downstream | |
|---|---|---|---|
| quasimodo (this study) | GGTAGTATTCGTAAGTTTGT | ACAATTCCAGCTCAACTCCC | seq confirmed |
| idefix (known/this study) | GGAAGTATCTGTAAGTTTAT | ATAATTCCAGACCAACCAAA | seq confirmed |
| 297 (this study) | CACCTGGAAGGTAACCAATC | AATTTCACAGGCACCTGGTA | seq confirmed |
| rover (this study) | GTTTTAATGAGTAAGTTAGA | CATTTTACAGGCCCCTTGTT | seq confirmed |
| 17.6 (this study) | ACACTTACTAGTAAGCTTGA | AATTTCACAGGCACCTGGCA | seq confirmed |

| **Springer subclade** | upstream | downstream | |
|---|---|---|---|
| active | | | |
| gtwin (this study) | TACAAGCCATGTAAGTTTGA | TGGTTTGCAGGTTCGCCCTC | seq confirmed |
| gypsy (known, this study) | AGAAAAGCATGTAAGTTTTA | TAATTTCCAGGTTCACCCTC | seq confirmed |
| gypsy6 (this study) | TACAGAACATGTACATCTTC | AATTTCACAGGATCGGACCA | seq confirmed |
| springer (this study) | CCTGAATCATGTAAGTGGGA | CACGTTACAGGTCGCTCCCA | seq confirmed |
| inactive | | | |
| gypsy2 (this study) | CAAAAAAAATGTAAGTGGGC | TCACCCACAGGACCAATCTT | predicted/not deregulated |
| gypsy3 (this study) | CACAGATTATGTAAGTGAGC | CATATTACAGGTCGTTTCCA | predicted/not deregulated |
| gypsy4 (this study) | TGAAATACATGTAAGTGATT | TTTATTACAGGTTAGGATAT | predicted/not deregulated |
| gypsy10 (this study) | AATATAACATGTAAGTGAGA | TCGCACTTAGGTTAGGAAAC | predicted/not deregulated |

| **Beagle subclade** | upstream | downstream | |
|---|---|---|---|
| HMS Beagle2 (this study) | CAATTTTATTGTGAGACAAG | TTACTTTCAGGATACTTACT | seq confirmed |

**Fig. S2: Conservation of *env-F* splice junctions.** Shown are sequence alignments of the experimentally identified *env-F* splice donor and acceptor junctions and their local conservation within the different subclades (intronic nucleotides in red are conserved in at least 80% of all sequences).

Supplementary Figure 3

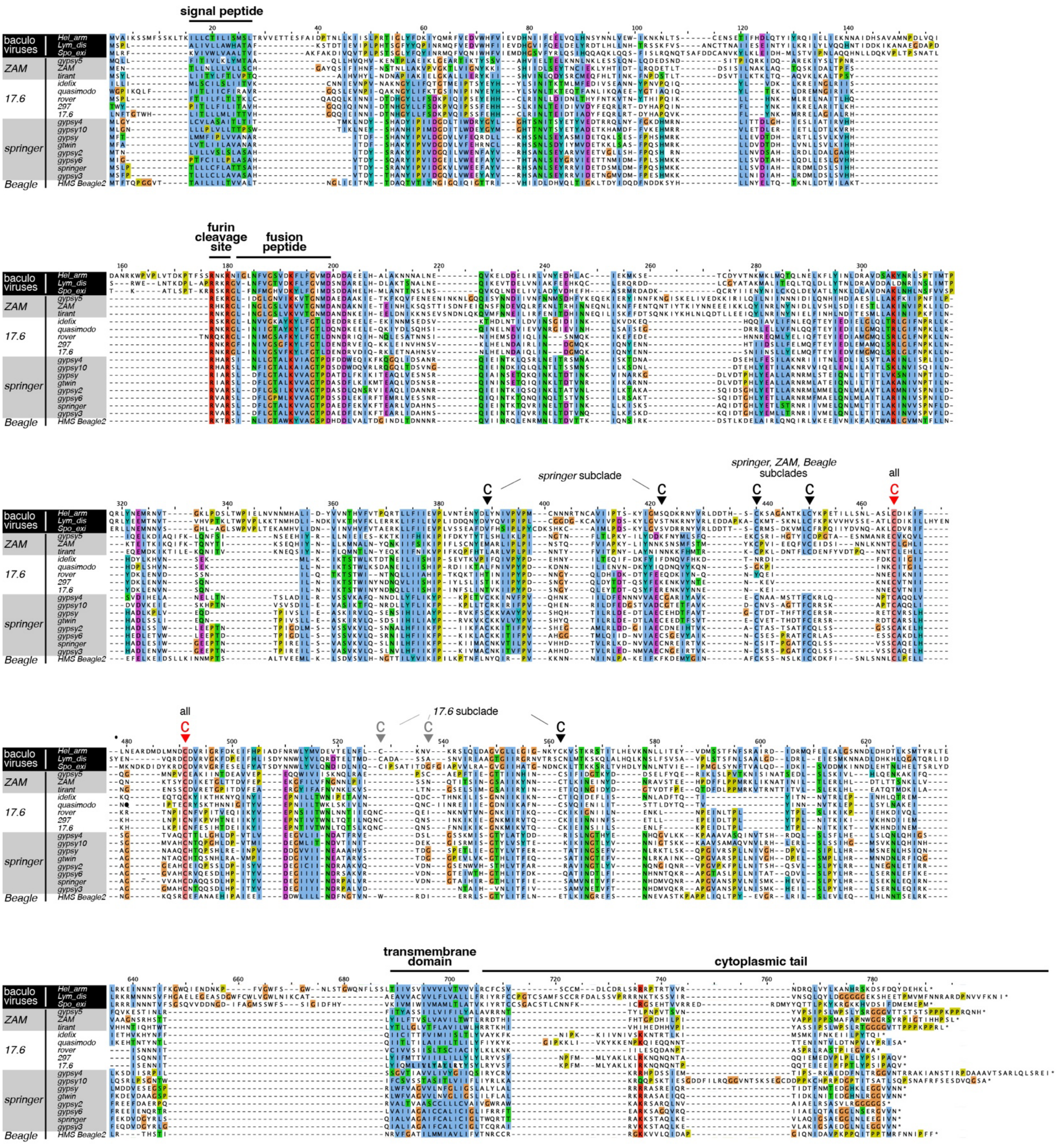

**Fig. S3: iERV Env-F proteins are related to F-type fusion glycoproteins from Baculoviruses.** Shown is a protein sequence alignment of all *Drosophila melanogaster* iERV spliced Envelopes, together with three F-type fusion glycoproteins from Baculoviruses isolated from indicated lepidopteran species. Indicated are N-terminal signal peptides, furin cleavage sites, fusion peptides, conserved Cysteine residues presumably involved in di-sulfide bond formation, transmembrane domains, and C-terminal cytoplasmic tails.

Supplementary Figure 4

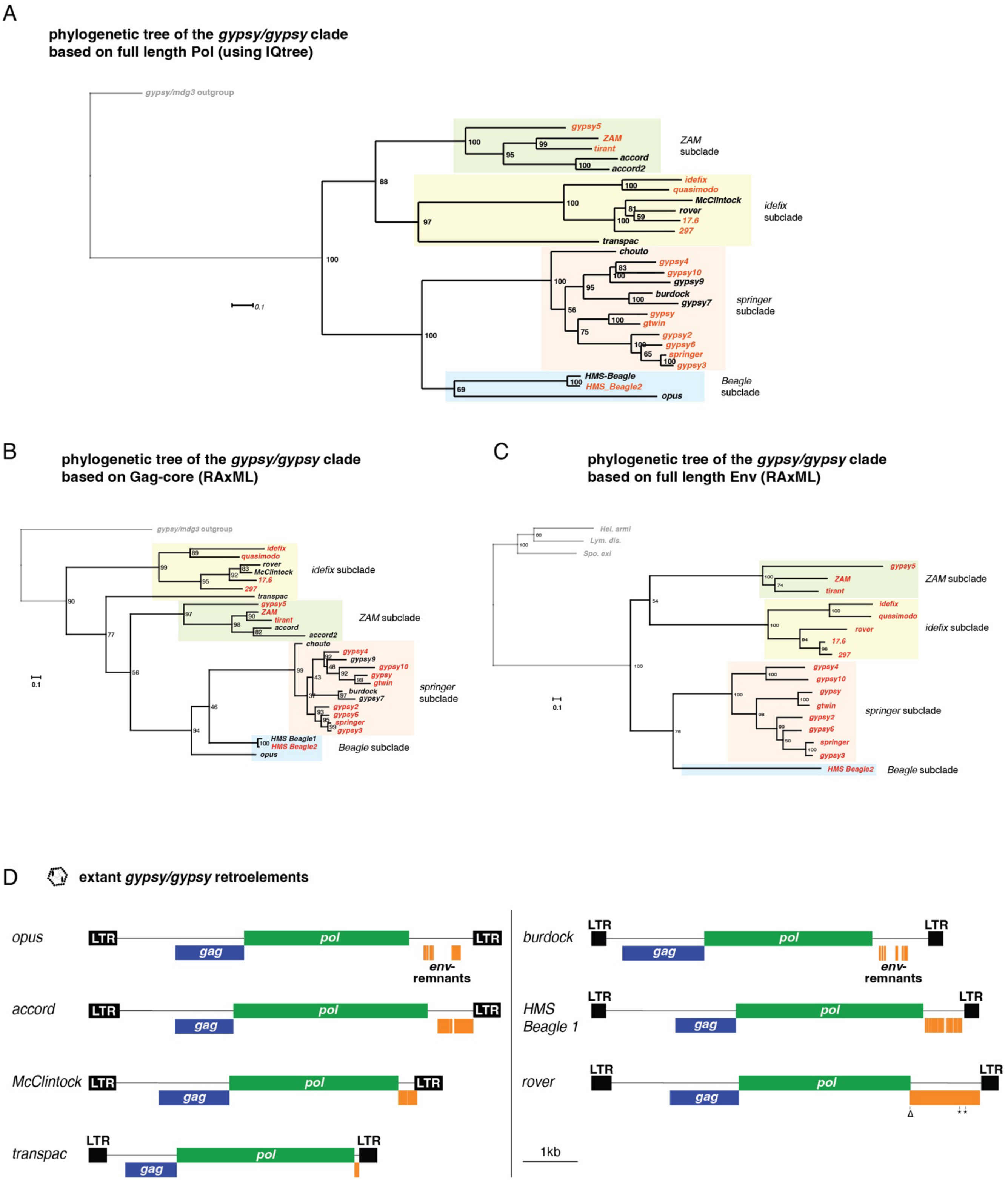

**Fig. S4: Phylogenetic trees for iERV Gag and Pol proteins. (A)** Shown are IQ-TREE estimated phylogenetic trees from the full length iERV Pol alignment with *gypsy/mdg3* outgroup as in Fig. 1C. Node labels reflect 1000 ultrafast bootstraps. Lineages with full length *env-F* are shown in red, those with mutated/lost *env-F* in black; subclades are color shaded. **(B-C)** Shown are RAxML estimated phylogenetic trees based on Gag-core domain and full length Env protein alignments from all iERV lineages (respective outgroups in grey). labelling as in (A). Note: Overall congruence of tree architecture with (A-C) and RAxML estimated Pol tree in Fig. 1C, despite the fact that Gag core domain diverged more than Pol and Env during evolution (see also Fig. 4A, Suppl. Fig. 14 A). **(D)** Cartoons (drawn to scale) of active iERV retroelements with identified fragments corresponding to *env-F* coding sequences indicated in orange (*env-F* remnants). For details on the *env-F* fragments see Suppl. Fig. 5.

#### Supplementary Figure 5

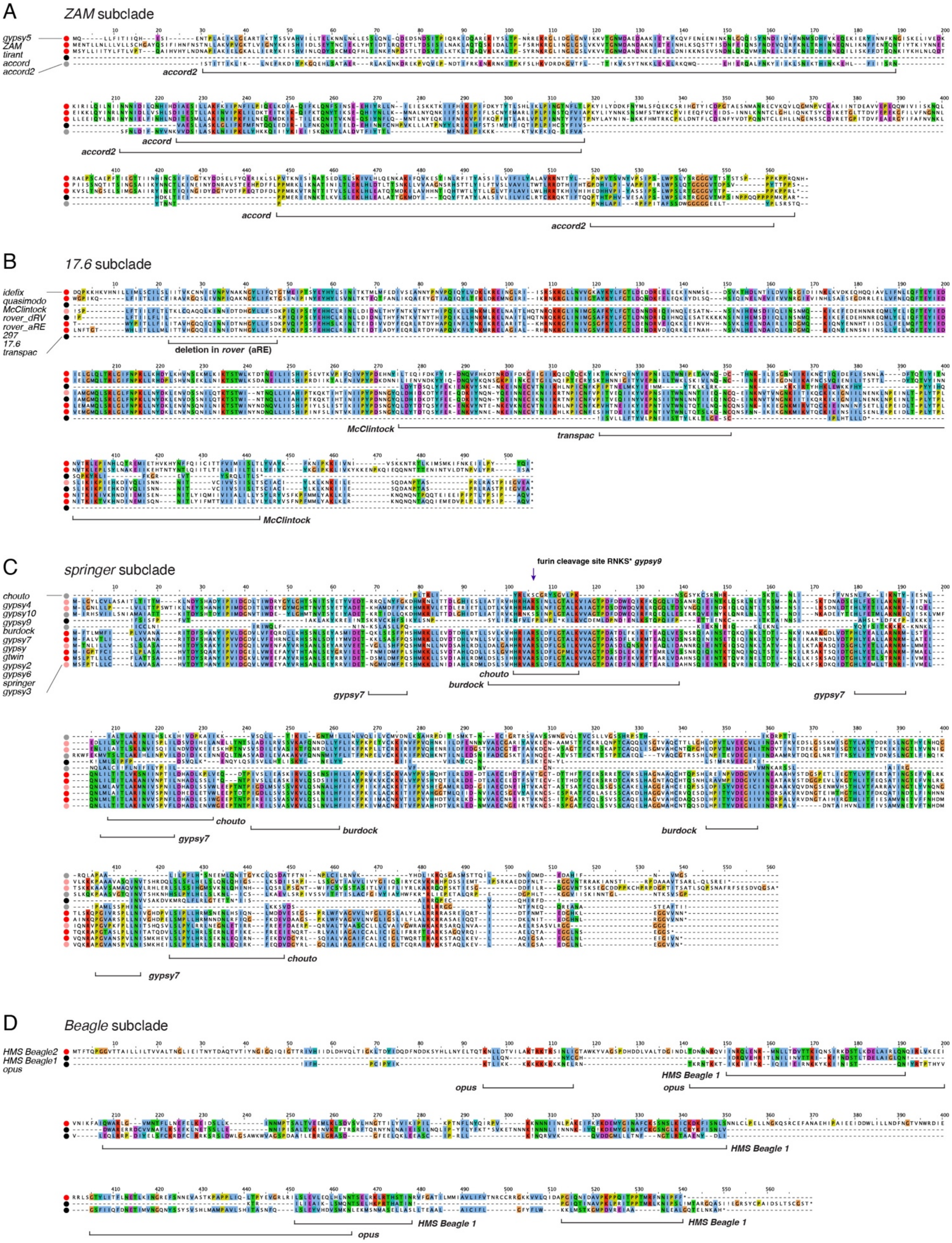

**Fig. S5: Sequence remnants of *env-F* in retroelement revertants among iERVs.** MACSE protein sequence alignments of Env-F and Env-F remnants within the individual iERV subclades. Dots to the left classify species as active retroviruses (dark red), inactive retroviruses (light red), active retroelements (black) and inactive retroelements (grey). Bracketed regions indicate *env-F* remnants present in derived LTR-retroelements lacking full length *env-F* genes.

Supplementary Figure 6

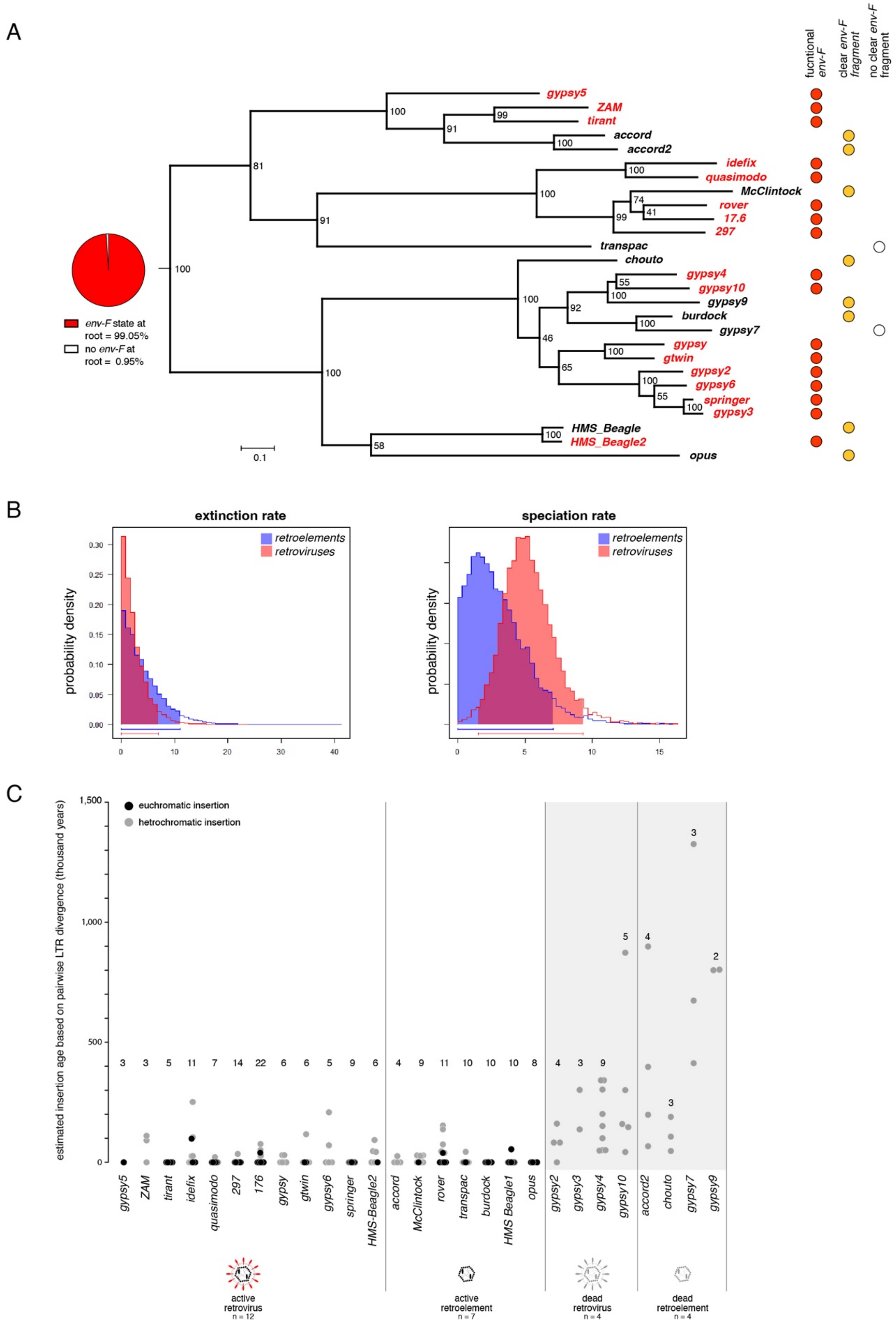

**Fig. S6: Ancestral character, diversification, and estimated age of iERV insertions in the *D. melanogaster* genome. (A)** Shown is the Pol alignment-based phylogenetic tree (RAxML) without outgroup and terminal branch associated *env-F* character states. Ancestral character reconstruction using Diversitree (BiSSE) estimates an *env-F*-harboring state at the root of the iERV clade with 99.1% probability. **(B)** BiSSE based Bayesian estimated differences of extinction (left panel) and speciation rates (right panel) for retroviruses (functional *env-F*) and retro-elements (no functional *env-F*) based on the tree in (A). Estimated speciation rates are higher for retroviruses, yet not statistically significant. Estimated extinction rates are not different. **(C)** Shown are estimated insertion ages (in thousands of years ago; TYA) calculated based on pairwise LTR divergence of all analyzed iERV insertions grouped into active and inactive retroviruses, and active and inactive retroelements (numbers above indicate the number of analyzed insertions).

Supplementary Figure 7

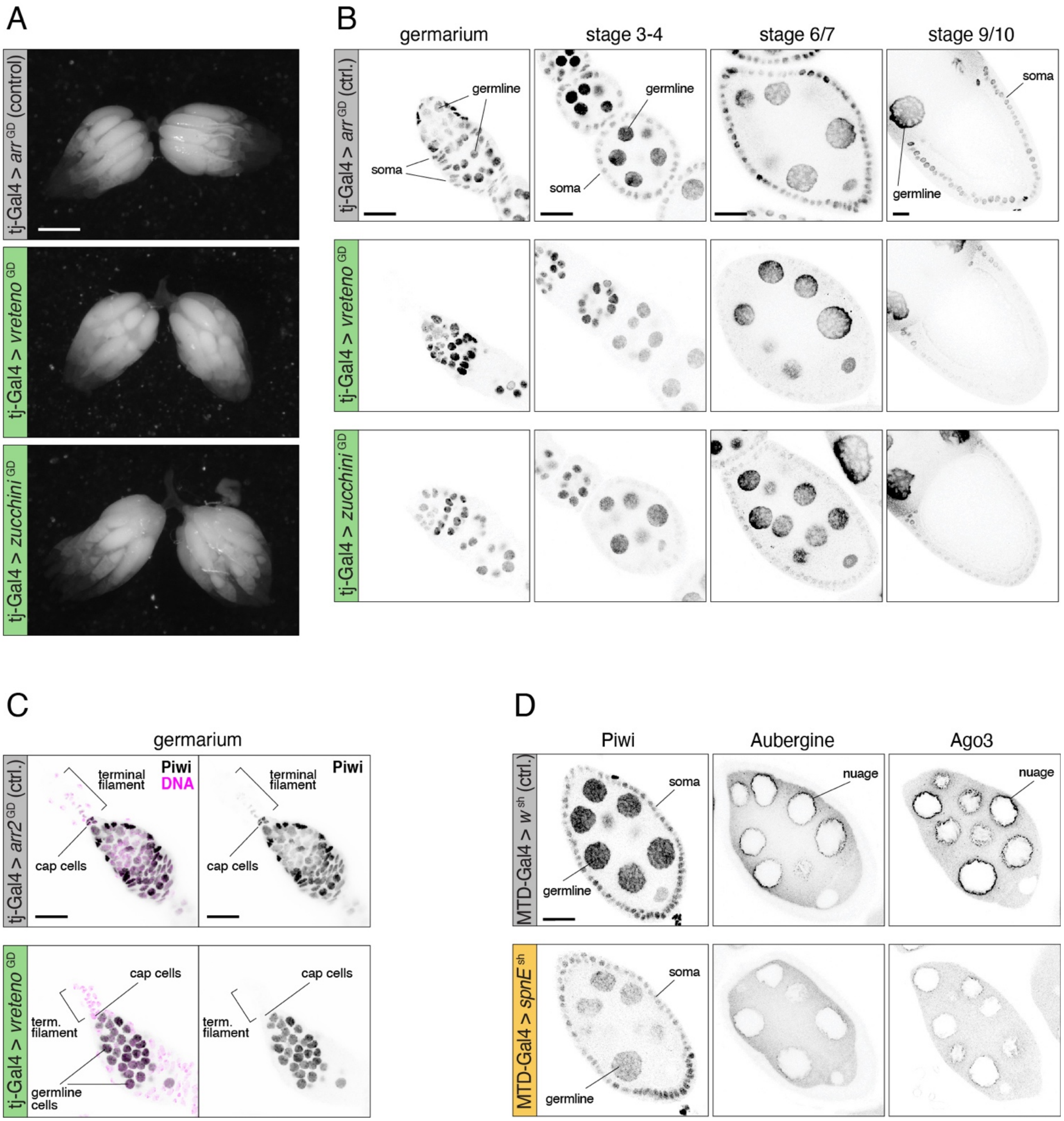

**Fig. S7: Validation of piRNA pathway knockdowns in ovarian soma and germline. (A)** Overall ovary morphology is not distorted upon piRNA pathway knockdowns in the soma (*tj*-Gal4 driven knockdown against *vreteno* or *zucchini* compared to control knockdown against *arrestin2*). **(B)** Antibody staining against Piwi (black) in ovaries (developmental stages indicated above) expressing long dsRNA hairpin constructs against *vreteno* or *zucchini* under *tj*-Gal4 control compared to control ovaries (Piwi protein levels are strongly reduced, specifically in somatic cells; scale bar: 20µm). **(C)** As in (B) but with a focus on terminal filament cells at the anterior tip of the germarium, which express low but detectable Piwi levels in a wildtype ovary, but no detectable Piwi in ovaries with defective somatic piRNA pathway (scale bar: 20µm). **(D)** Shown are antibody stainings against Piwi, Aub, and Ago3 in control ovaries and in ovaries expressing a short hairpin RNA against *spn-E* specifically in germline cells under control of the MTD-Gal4 driver (scale bar: 20µm; for the *aub* and *aub*+*ago3* knockdown conditions, see (*43*)). Loss of nuage localization for Aub and Ago3 as well as reduced levels for nuclear Piwi are indicative of a strong disruption of the germline piRNA pathway.

#### Supplementary Figure 8

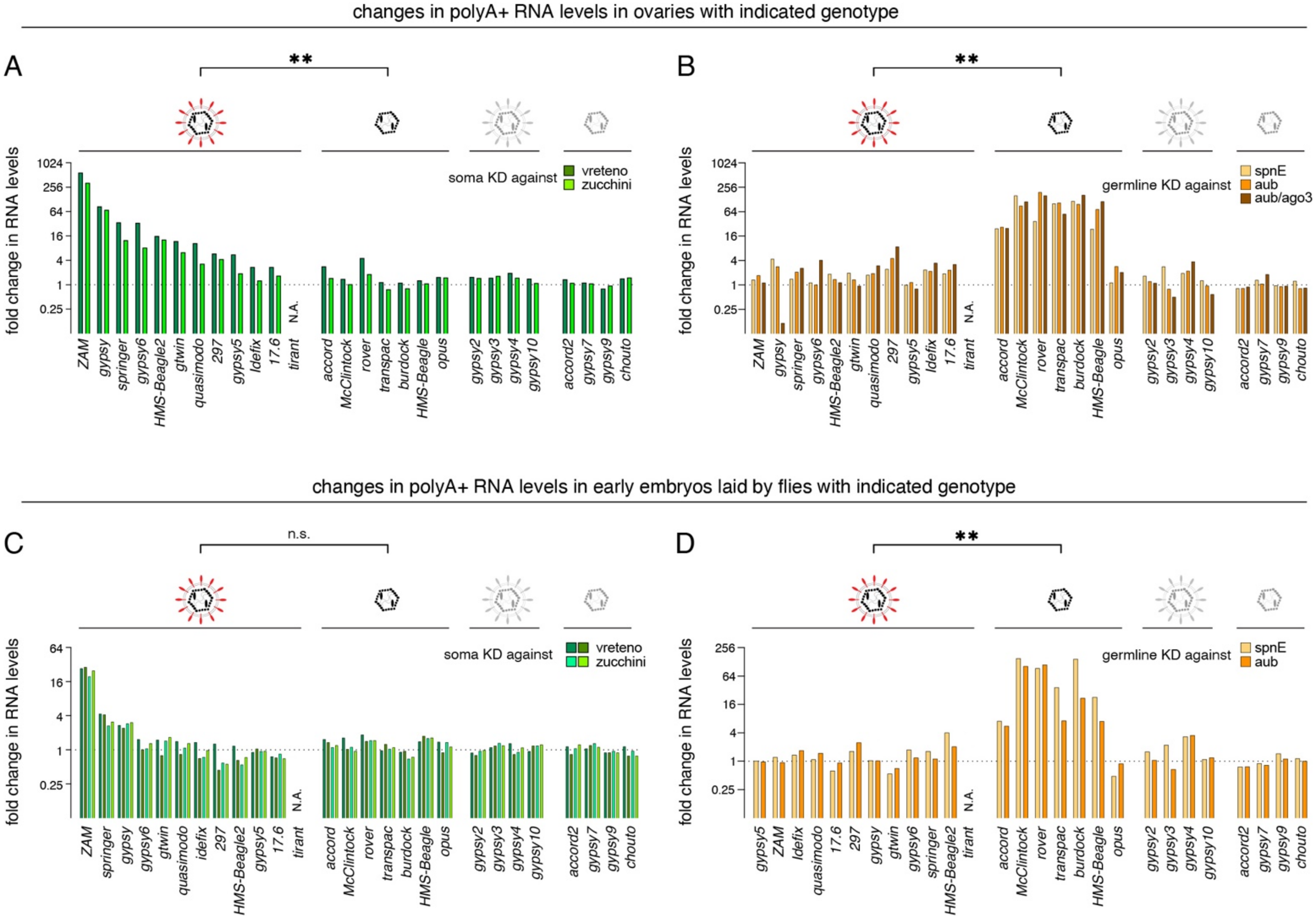


**Fig. S8: Changes in iERV transcript levels in ovaries and early embryos upon piRNA pathway loss. (A, B)** Shown are the fold-changes of all iERV poly-A$^{+}$ RNA levels in ovaries lacking the piRNA pathway in the soma (A) or in the germline (B) over control ovaries. **(C, D)** Shown are the fold-changes of all iERV poly-A$^{+}$ RNA levels in early embryos laid by flies lacking piRNA pathway control in the soma (C) or germline (D). The *tirant* retrovirus cannot be analyzed as it is not present in most of our experimental strains (N.A.) iERVs are grouped into active retroviruses, active retroelements, inactive retroviruses and inactive retroelements. Statistical significance (** indicates $p < 0.01$) was calculated according to Mann-Whitney (two-tailed test).

#### Supplementary Figure 9

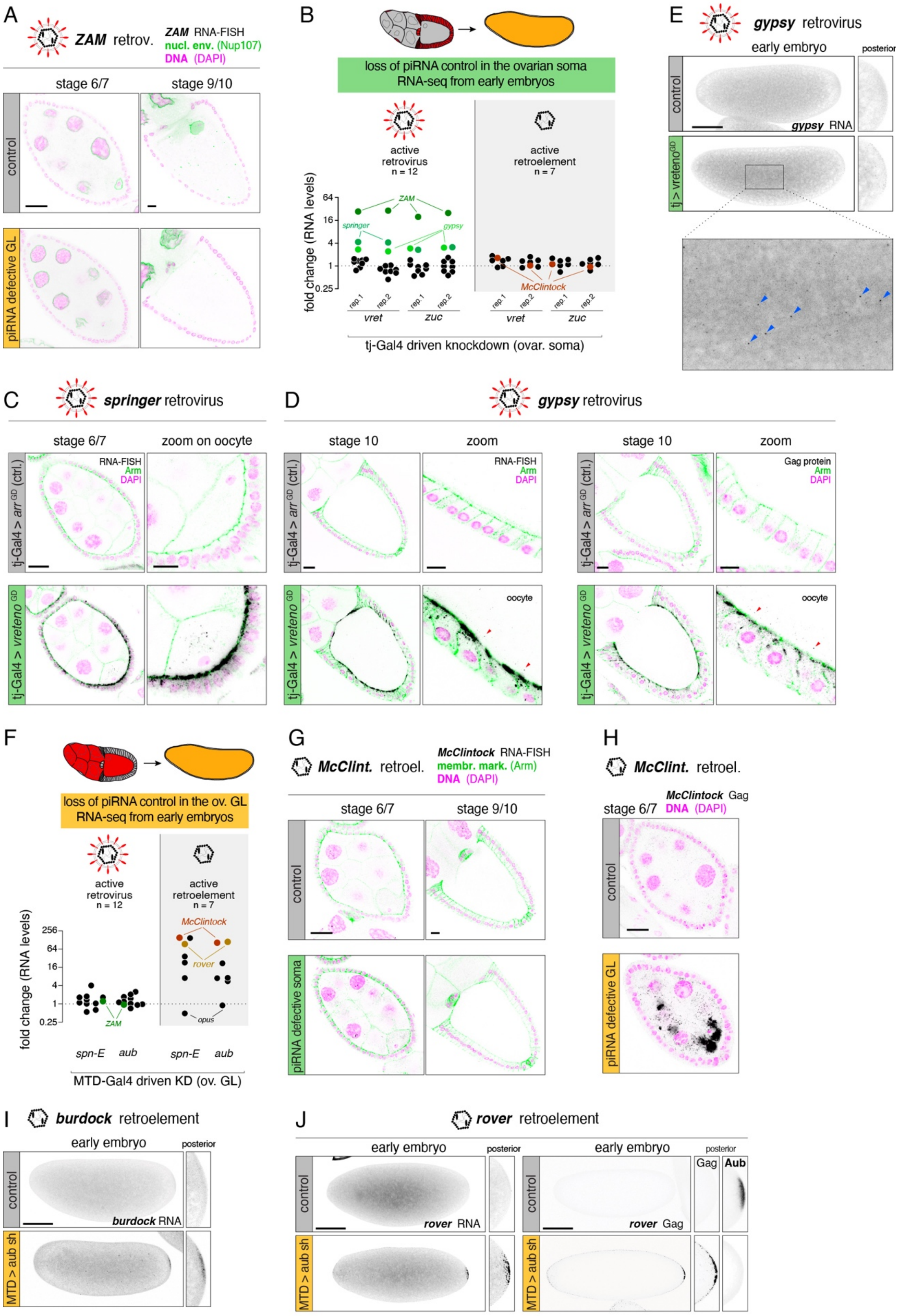

**Fig. S9: De-repression of iERVs in ovaries or early embryos upon loss of the piRNA pathway.** **(A)** Full repression of the retrovirus *ZAM*, judged by RNA smFISH, in control ovaries as well as in ovaries with defective germline piRNA pathway (genotypes are *MTD*-Gal4 driven shRNAs against *aub*+*ago3* or *white*; DAPI labels nuclei (magenta), *ZAM* RNA is in black, GFP-Nup107 labels nuclear envelopes in green; scale bars: 20µm). **(B)** Jitter plot indicating the fold changes in iERV steady state polyA$^+$ RNA levels in 0-60 min old, dechorionated embryos laid by mothers with defective somatic piRNA pathway versus those laid by control mothers (RNAi and GAL4 strains used for transgenic RNAi are indicated; replicates are biological replicates). **(C, D)** Detection of *springer* (D) or *gypsy* (E) transcripts by RNA smFISH or *gypsy* Gag protein (E) in egg chambers of indicated age and genotype. DAPI labels nuclei (magenta), anti-Armadillo staining labels cell outlines in green, red arrowheads point to RNA smFISH signal (black) in the developing oocyte, indicative of soma-to-germline transfer (scale bars: 20µm for full egg chambers and 10µm for the enlarged region). **(E)** Detection of *gypsy* transcripts by RNA-smFISH in pre-blastoderm embryos (<32 nuclei), laid by mothers with defective somatic piRNA pathway (maternal genotype: *tj*-Gal4 > *vreteno*$^{GD}$ or *arrestin2*$^{GD}$*;* scale bar: 100µm). Images represent maximum intensity Z projections of confocal stacks. Individual smFISH dots are shown in the zoom-in image below. **(F)** Jitter plots indicating the fold changes in iERV steady state polyA$^+$ RNA levels in 0-60 min old, dechorionated embryos laid by mothers with defective germline piRNA pathway versus those laid by control mothers (RNAi and GAL4 strains used for transgenic RNAi are indicated; replicates are biological replicates). **(G)** Lack of expression of the retroelement *McClintock*, judged by RNA smFISH, in control ovaries as well as in ovaries with defective somatic piRNA pathway (genotypes are *tj*-Gal4 driven dsRNA hairpins against *vreteno* or *arrestin2* as control; DAPI labels nuclei (magenta), *McClintock* RNA shown in black, anti-Armadillo labels cell outlines in green; scale bars: 20µm). **(H)** *McClintock* Gag in stage 6/7 egg chamber detected by immuno-fluorescence in ovaries with defective germline piRNA pathway (bottom) or control ovaries (top). Note the accumulation of *McClintock* capsid protein in the developing and transcriptionally inactive oocyte (scale bar: 20µm). **(I, J)** Detection of *burdock* (I), or *rover* (J) transcripts by RNA-smFISH in pre-blastoderm embryos (<32 nuclei), laid by mothers with defective germline piRNA pathway (maternal genotype: *MTD*-Gal4 driven shRNA against *aub* or *white;* scale bars: 100µm). In the case of *rover*, the same analysis was also done with an anti-*rover* Gag antibody and with an anti-Aub staining serving as knockdown control (J). Images represent maximum intensity Z projections of confocal stacks. The zoom-in images focus on the posterior pole where the future germline cells will form.

#### Supplementary Figure 10
page 1

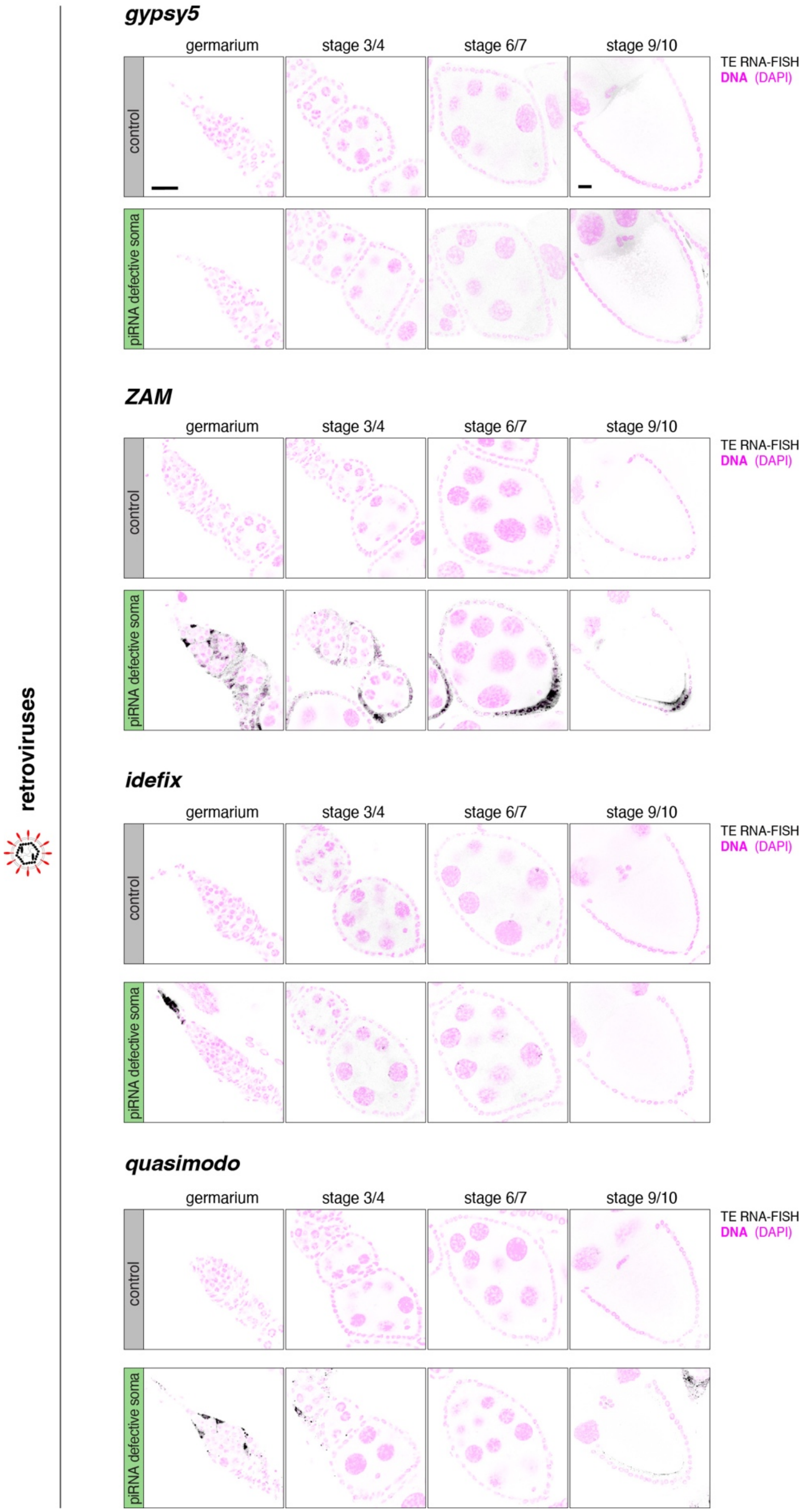

#### Supplementary Figure 10
page 2

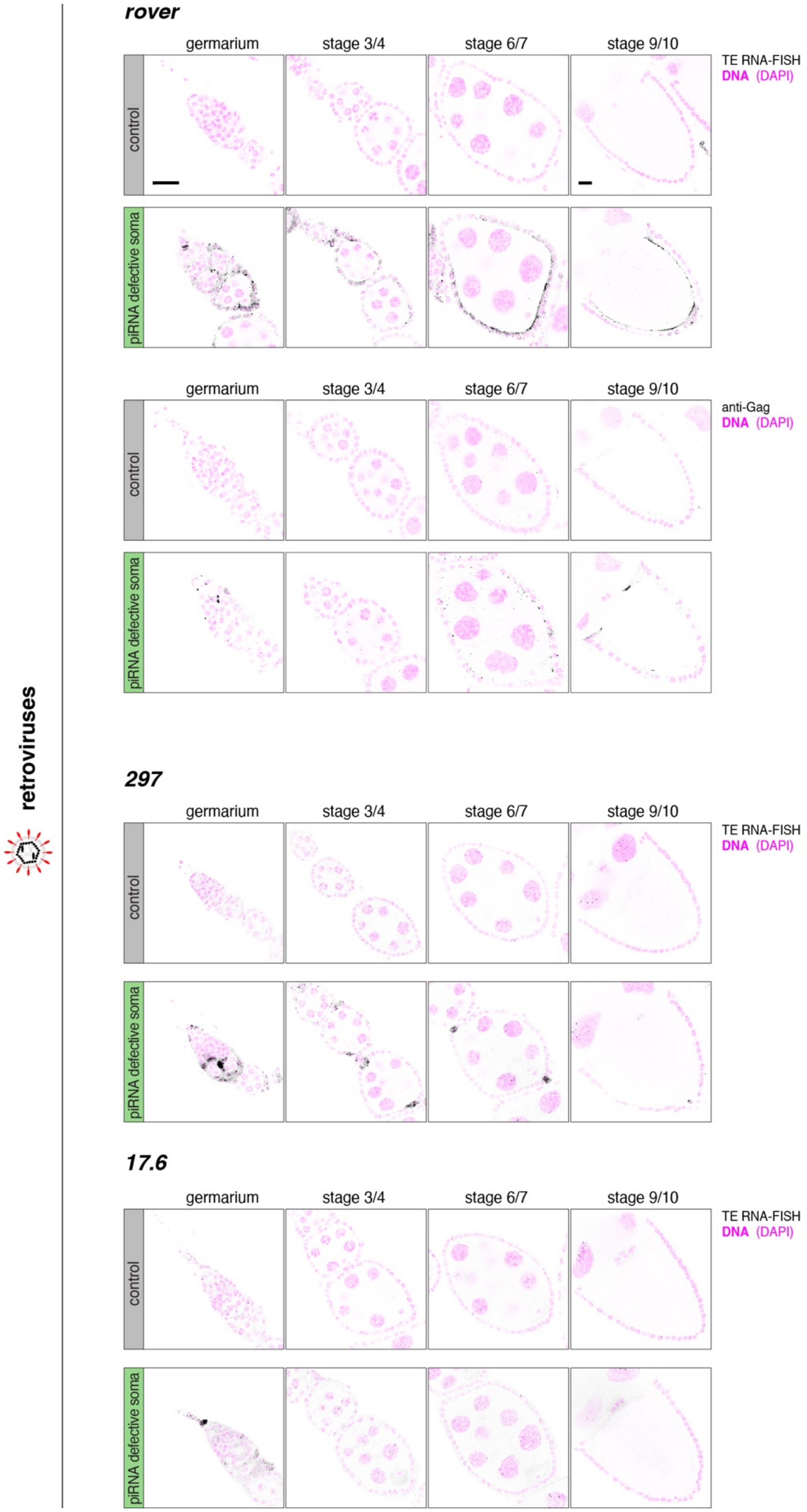

Supplementary Figure 10
page 3

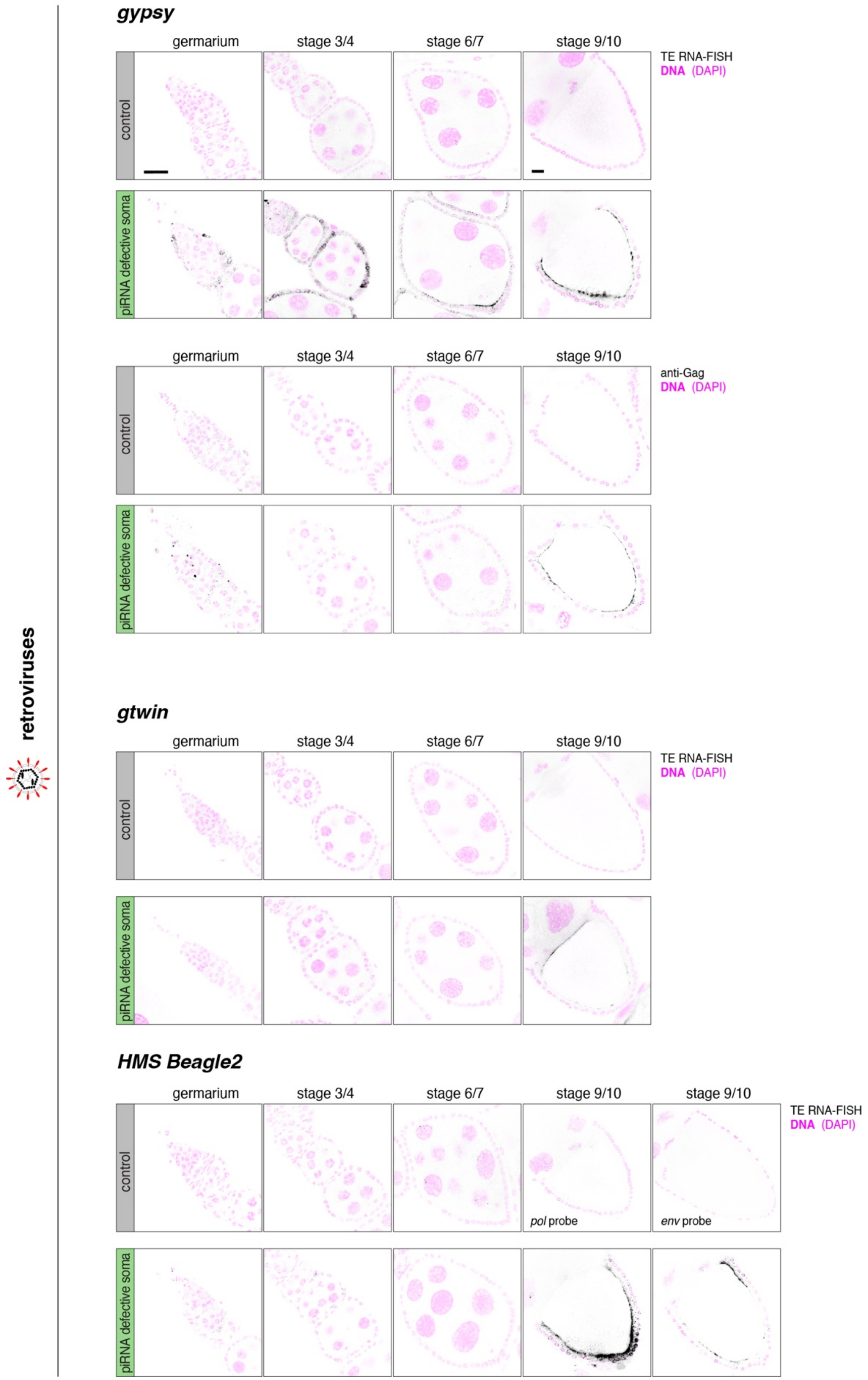

Supplementary Figure 10
page 4

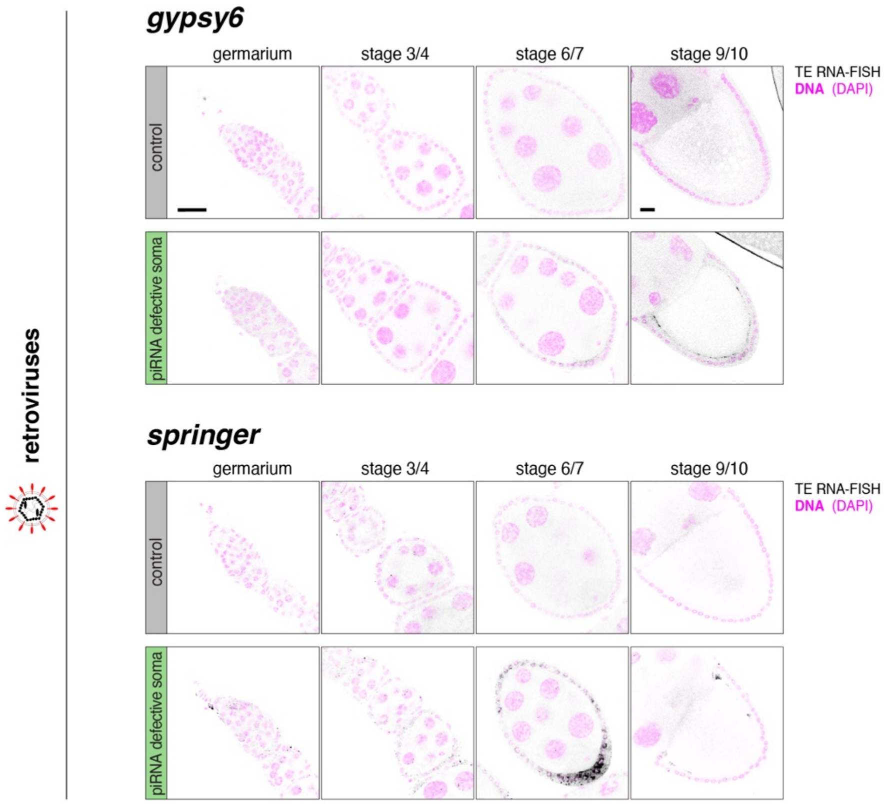


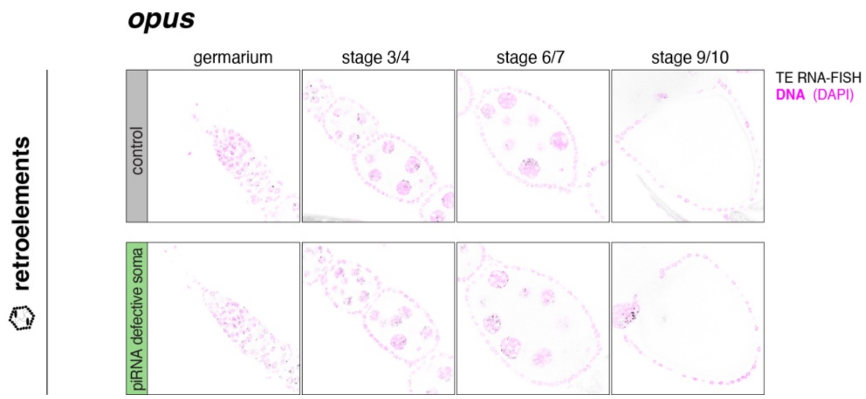

Supplementary Figure 10
page 5

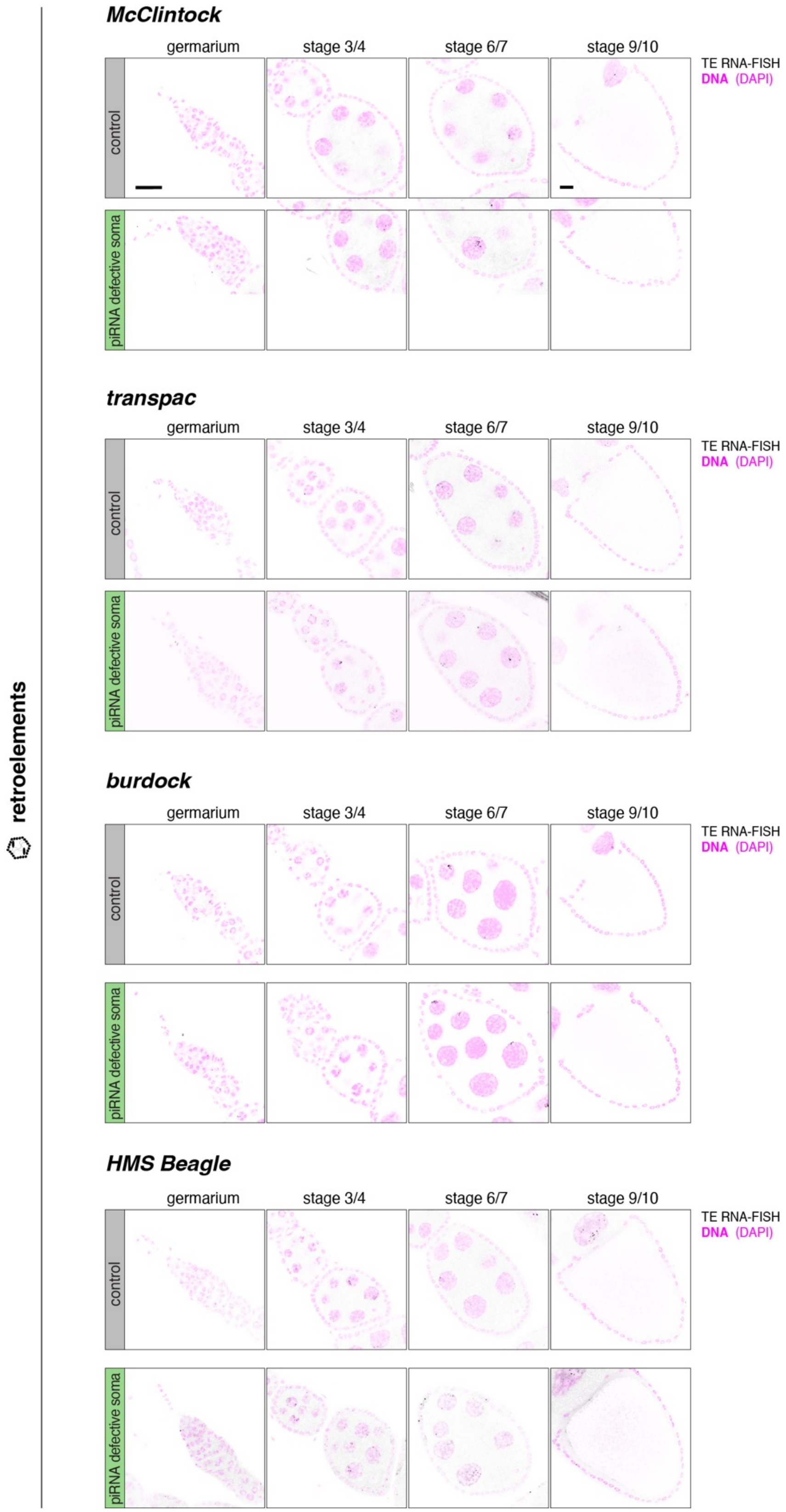

**Fig. S10: Systematic expression analysis of all active iERV lineages in ovaries lacking somatic piRNA pathway control.** Shown are smFISH experiments and immuno-fluorescence experiments against indicated iERVs (retroviruses and retroelements) in control ovaries and in ovaries with defective piRNA pathway in the soma. All stages of oogenesis are shown, scalebars equal 20µm.

Supplementary Figure 11

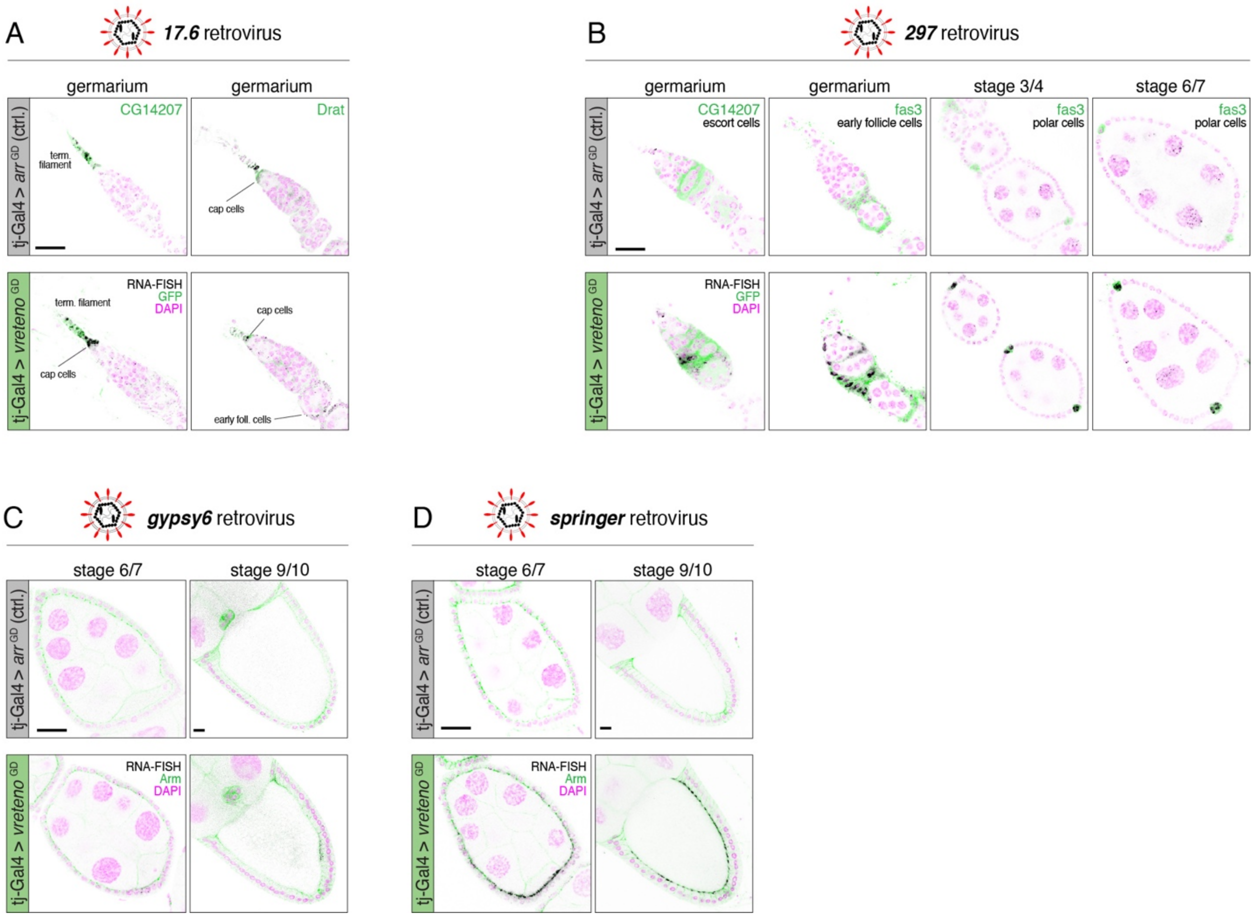


**Fig. S11: Niche expression of infectious iERVs in the ovarian soma.** Detection of indicated iERV transcripts by RNA-smFISH (black) in egg chambers of indicated oogenesis stages and genotypes. Panel **(A)** shows *17.6*, **(B)** *297*, **(C)** *gypsy6*, and **(D)** *springer*; DAPI is shown in magenta, indicated GFP-traps (A, B) or anti-Armadillo staining (C-D) in green; scale bars: 20µm.

Supplementary Figure12

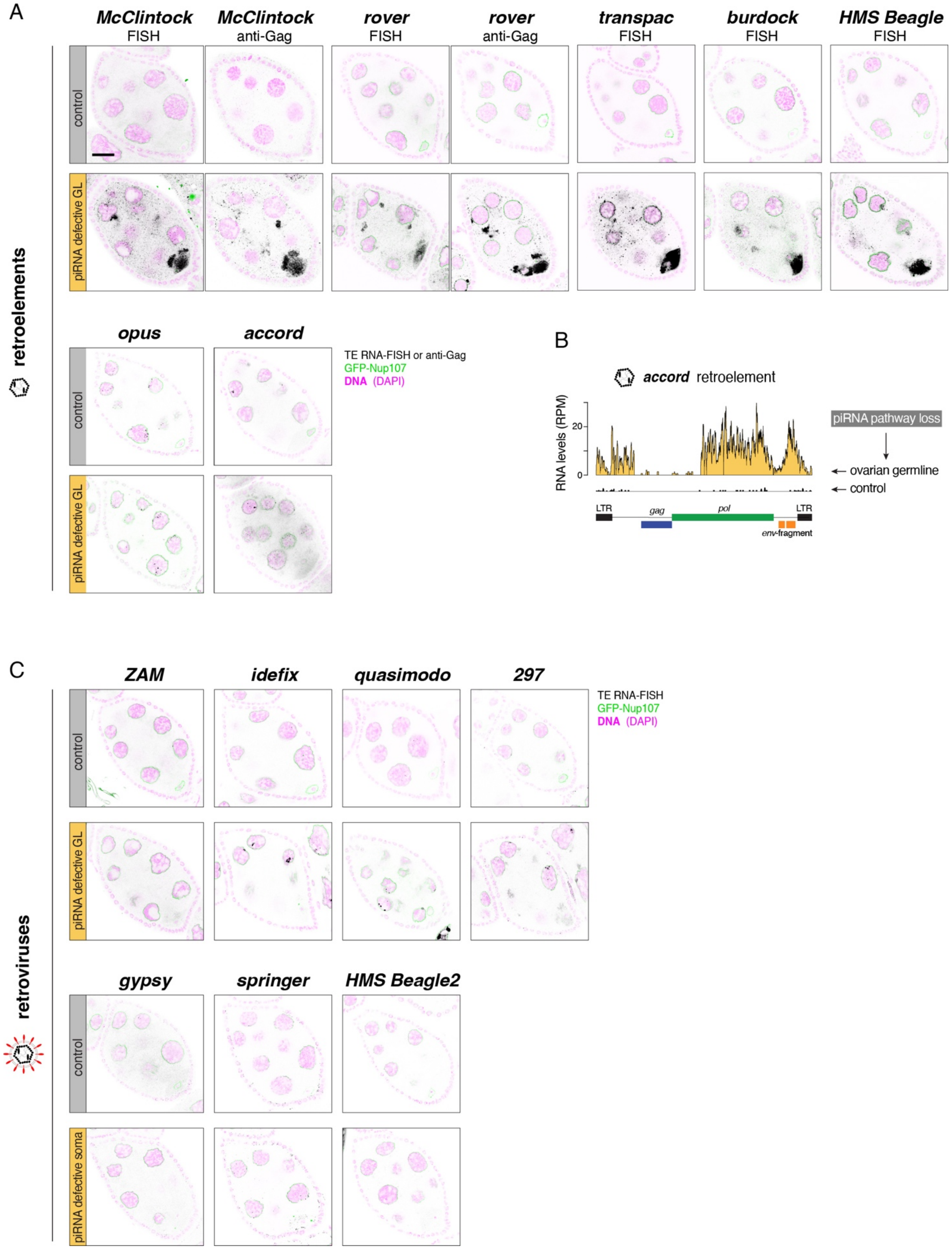

**Fig. S12: Systematic expression analysis of all active iERV lineages in ovaries lacking germline piRNA pathway control. (A)** Shown are smFISH experiments and immuno-fluorescence experiments against indicated iERVs (retroelements) in control ovaries and in ovaries with defective piRNA pathway in the germline. Shown are only stage 6/7 egg chambers as other stages did not add additional insight. Scalebars equal 20µm. **(B)** Shown is the increase in *accord* RNA levels as measured by poly-$A^+$ RNA-seq in ovaries lacking piRNA pathway control in the germline. Due to the gap in read coverage, we suspect that our experimental flies lack a full-length *accord* insertion. **(C)** As in (A) but for retroviruses.

### Supplementary Figure 13

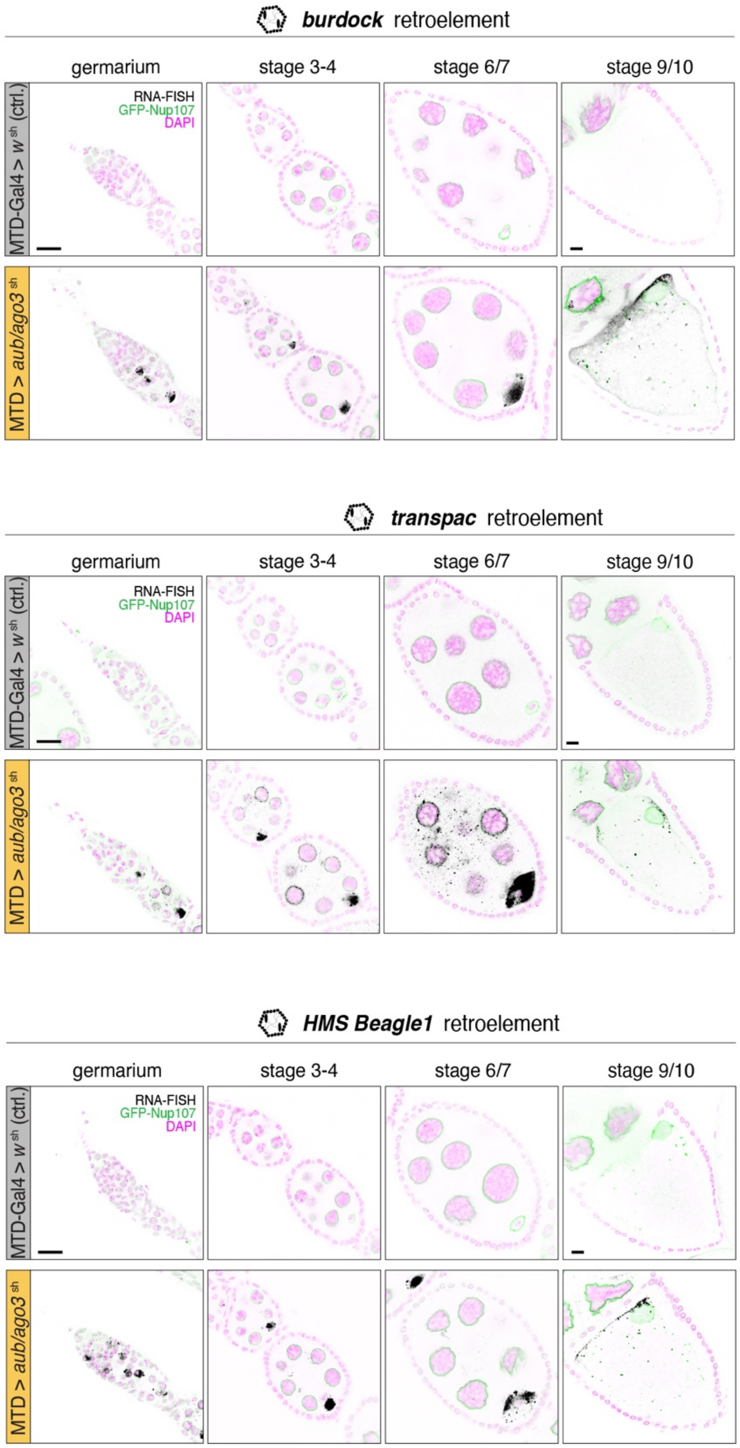

**Fig. S13: Germline specific expression of non-infectious iERVs in the ovarian germline.** Detection of indicated iERV transcripts by RNA-smFISH in germaria or egg chambers of indicated oogenesis stages and genotypes (DAPI labels nuclei (magenta), GFP-Nup107 labels nuclear envelopes in green, and smFISH signal is shown in black; scale bars: 20µm). Retroelement transcripts start to be detectable in differentiating germline cystoblasts in the germarium and are enriched in the developing oocyte where they accumulate around the oocyte nucleus.

Supplementary Figure 14

A

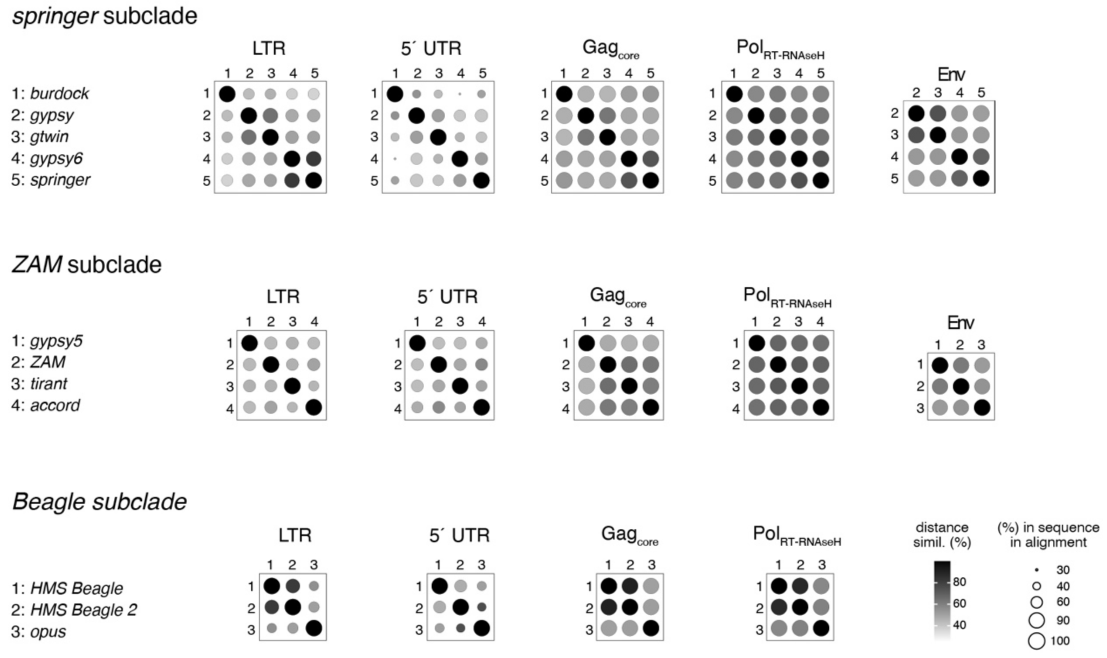


B

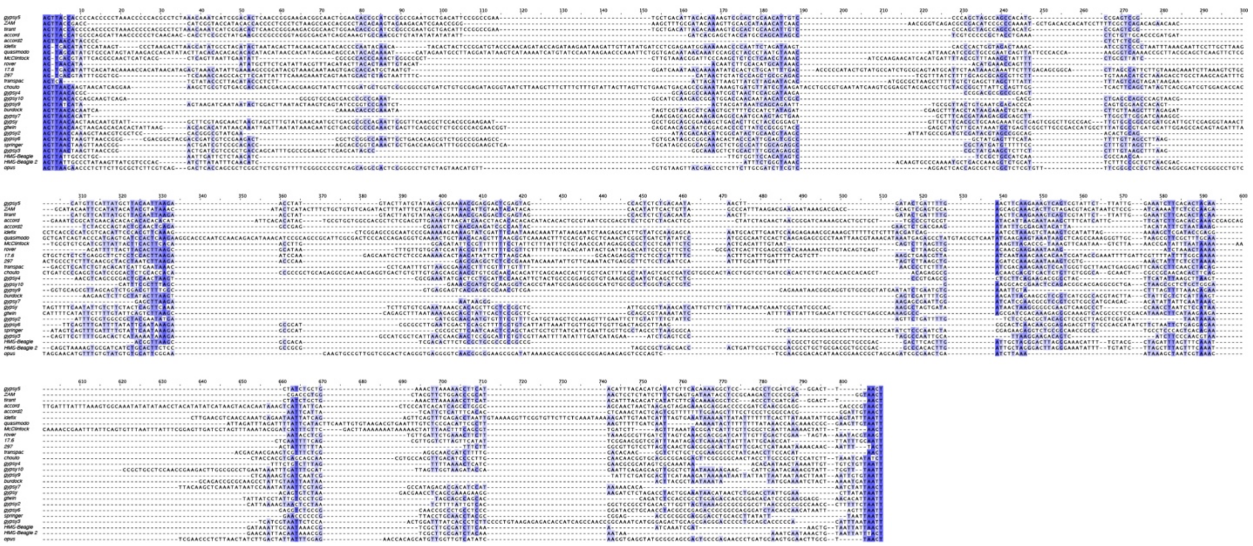

**Fig. S14: LTR and 5´ UTR sequence divergence among iERVs. (A)** Shown are pairwise similarities between the DNA sequences for LTR, 5´ UTR, the protein domains Gag core, $Pol_{RT-RNAseH}$, and full length spliced Env-F. Circle size indicates percentage of nucleotides in aligned positions, greyscale indicates pairwise similarities among iERVs, separately for the indicated subclades (see Fig. 4 for the *idefix* subclade). **(B)** DNA sequence alignment of all iERV LTR sequences.

Supplementary Figure 15

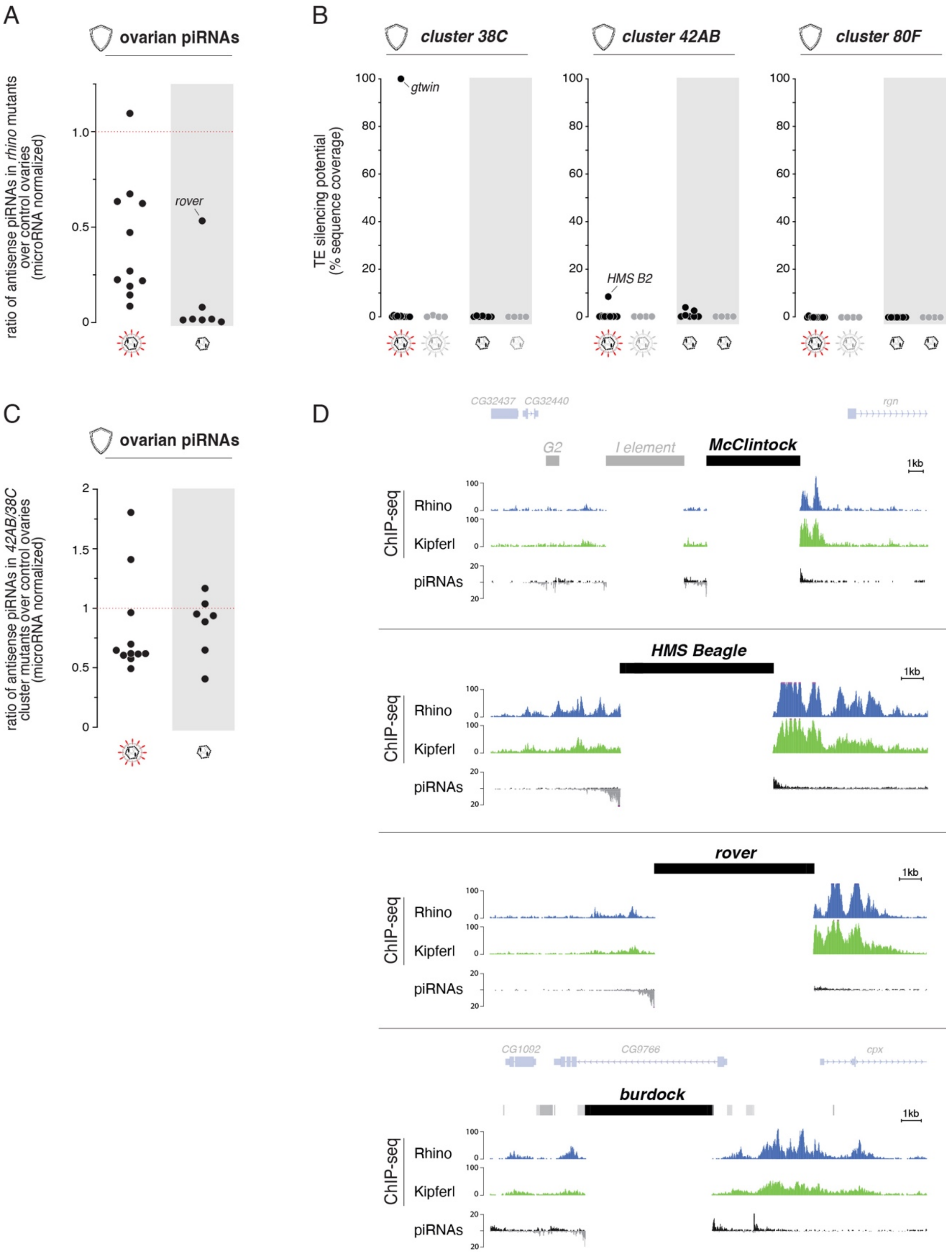

**Fig. S15: piRNAs targeting retroelements in the germline are Rhino-dependent. (A)** Shown are ratios of antisense piRNAs (microRNA normalized) mapping to active retroviruses (left) or retroelements (right) among iERVs from ovaries of germline specific *rhino* RNAi flies (*MTD*-Gal4 driven shRNA against *rhino*) versus control ovaries (*MTD*-Gal4 driven shRNA against *white*). The transition element *rover* is the only retroelement with Rhino-independent piRNAs, in line with an almost full-length *rover* insertion in *flamenco*. **(B)** Shown are the silencing potentials of dual-strand germline piRNA clusters against active and inactive retroviruses and retroelements among the iERVs. Silencing potentials are represented as percent coverage of TE sequences found within the respective clusters (calculated as 25mers with zero mismatches). **(C)** Shown are ratios of antisense piRNAs (microRNA normalized) mapping to active retroviruses (left) or retroelements (right) from ovaries of *cluster 42AB/38C* double-mutant flies versus control ovaries. **(D)** Shown are stand-alone insertions of the indicated retroelements that act as mini-piRNA source loci in the germline of *iso-1*. Genome-unique reads from Rhino ChIP-seq, Kipferl ChIP-seq and ovarian piRNAs (all from the *iso-1* strain) are shown. Reads mapping to the indicated TE insertions are multi-mapping and therefore not displayed.

Supplementary Figure 16

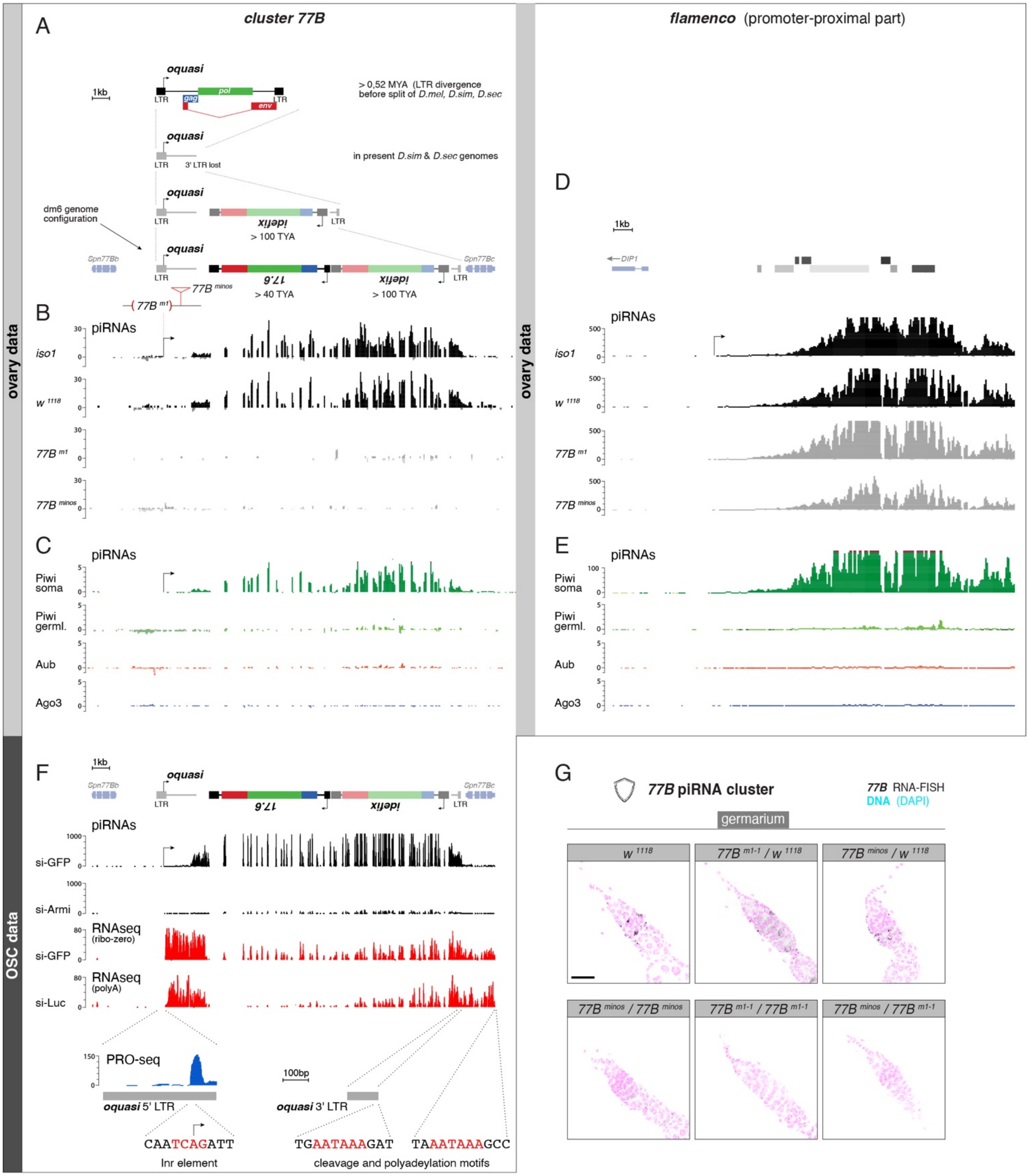

**Fig. S16: Characterization of the somatic *77B* piRNA cluster. (A)** Cartoon depicting the proposed evolutionary trajectory of the *77B* piRNA cluster that originated from an old *quasimodo* retrovirus (*oquasi*) insertion between the *Spn77Bb* and *Spn77Bc* genes and captured an *idefix* and *17.6* retroviral insertion in antisense orientation (scale and alignment of the cartoon corresponds to the data in panels B, C, F). **(B, C)** Shown are genome-unique piRNA mappings from ovaries of indicated genotypes (B) or bound to indicated PIWI-clade proteins from wildtype ovaries (C) (*43*). Panel C shows that *77B*-derived piRNAs are specifically bound to soma-expressed Piwi). **(D, E)** As in panels B and C, but for the beginning of the *flamenco* piRNA cluster. **(F)** Analysis of *cluster 77B* expression in ovarian somatic cells (OSCs), a cell line derived from ovarian follicle stem cells. Shown are genome-unique mappings of piRNA reads, RNA-seq reads (from ribo-zero and from poly A-plus libraries), and PRO-seq reads (all mappers) at the *77B* piRNA cluster. At the bottom, motifs for transcription initiation (Initiator element) and cleavage/poly-adenylation are shown. **(G)** Expression analysis of the *77B* piRNA cluster in the germarium of ovaries with indicated genotype based on RNA-smFISH (black) (DAPI labels nuclei (magenta); images represent maximum intensity projections of ten confocal Z sections).

Supplementary Figure 17

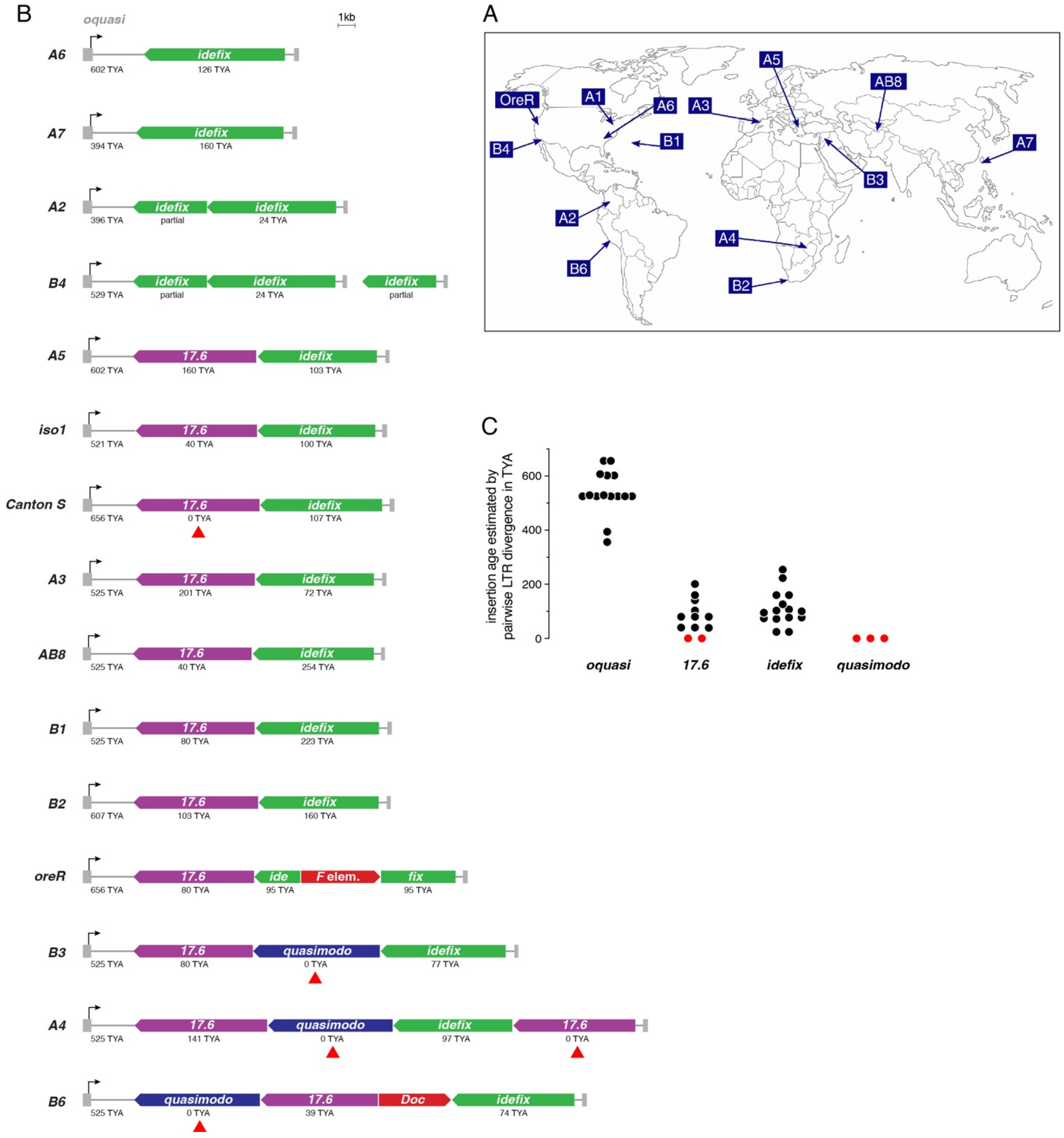

**Fig. S17: Sequence variations in the somatic *77B* piRNA cluster in DSPR strains. (A)** Earth map indicating the collection sites for the *Drosophila melanogaster* DSPR strains. Image unmodified from (*64*). **(B)** Sequence content of the *77B* piRNA cluster in all DSPR strains. Below each annotated LTR element the estimated age of the insertion, based on pairwise LTR divergence, is indicated (TYA: thousand years ago). Red arrowheads mark recent insertions (modern *quasimodo* and *17.6*) with no LTR divergence, suggesting ongoing cluster evolution. **(C)** Jitter plot summarizing the age of the *oquasi*, *17.6*, *idefix*, and *quasimodo* insertions within the *77B* piRNA cluster of all analyzed DSPR strains. Red dots mark recent insertions (modern *quasimodo* and *17.6*) with no LTR divergence.

Supplementary Figure 18

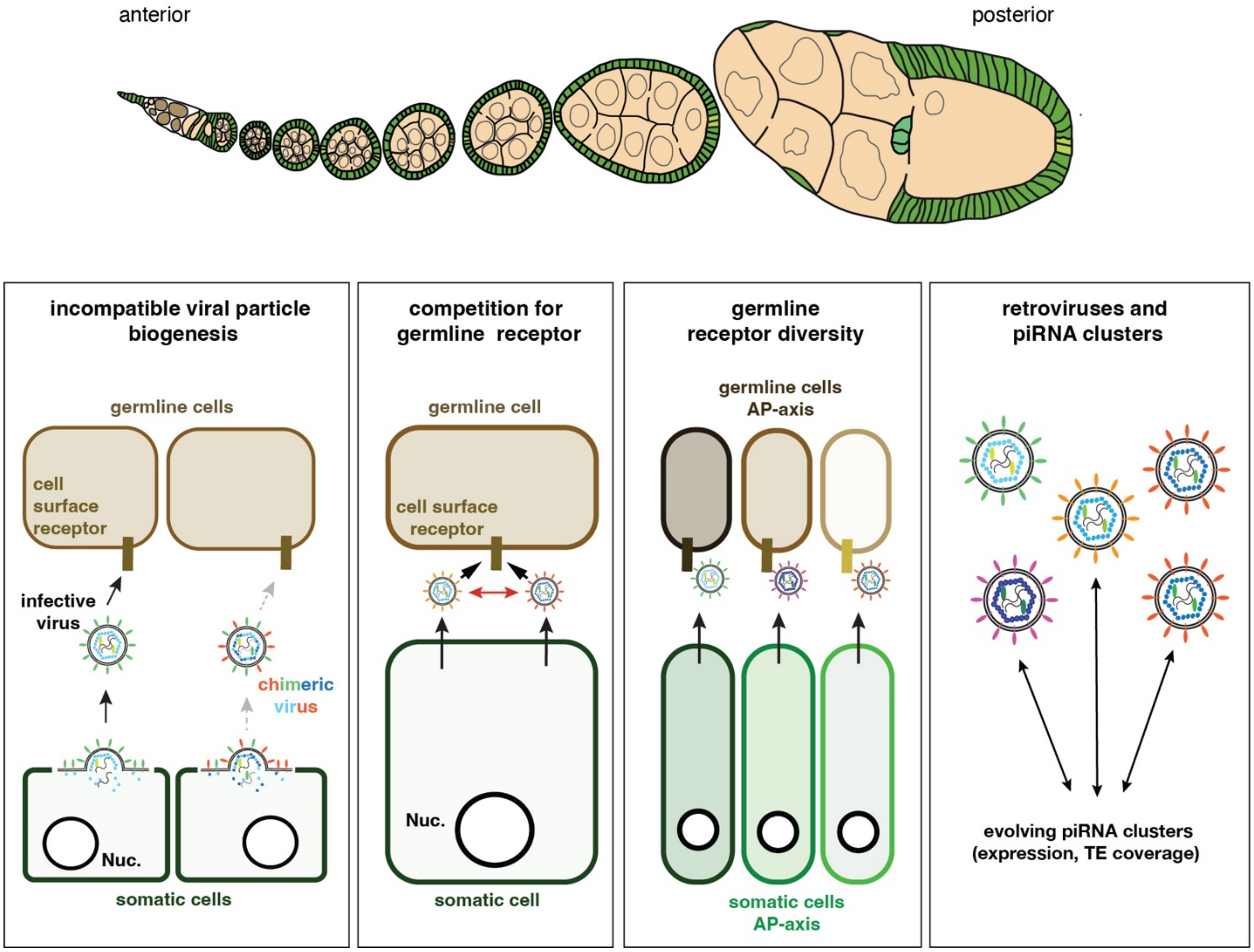


**Fig. S18: Potential scenarios of competition between infectious iERVs and between iERVs and the *Drosophila* host.**
